# DNA Tetrahedral Nanocages as a Promising Nanocarrier for Dopamine Delivery in Neurological Disorders

**DOI:** 10.1101/2023.09.19.558434

**Authors:** Ramesh Singh, Krupa Kansara, Sandip Mandal, Ritu Varshney, Sharad Gupt, Ashutosh Kumar, Prabal K Maiti, Dhiraj Bhatia

## Abstract

Dopamine is a neurotransmitter in the central nervous system that is essential for many bodily and mental processes, and a lack of it can cause Parkinson’s disease. DNA tetrahedral (TD) nanocages are promising in bio-nanotechnology, especially as a nanocarrier. TD is highly programmable, biocompatible, and capable of cell differentiation and proliferation. It also has tissue and blood-brain barrier permeability, making it a powerful tool that could overcome potential barriers in treating neurological disorders. In this study, we used DNA-TD as a carrier for Dopamine to cells and zebrafish embryos. We investigated the mechanism of complexation between TD and dopamine hydrochloride using gel electrophoresis, fluorescence and circular dichroism (CD) spectroscopy, atomic force microscopy (AFM), and molecular dynamic (MD) simulation tools. Further, we demonstrate these Dopamine-loaded DNA tetrahedral nanostructures’ cellular uptake and differentiation ability in SH-SY5Y neuroblastoma cells. Furthermore, we extended the study to zebrafish embryos as a model organism to examine survival and uptake. The research provides valuable insights into the complexation mechanism and cellular uptake of dopamine-loaded DNA tetrahedral nanostructures, paving the way for further advancements in nanomedicine for Parkinson’s disease and other neurological disorders.

## Introduction

Neurotransmitters are endogenous chemical messengers that play a crucial role in the complex network of neurons in the body. They facilitate neurons to communicate and transmit signals from neurons to targeted cells.^1,2^ An imbalance in the physiological concentration of specific neurotransmitters causes neurological disorders such as Parkinson’s disease (PD), Alzheimer’s disease, etc. Dopamine is a critical neurotransmitter in the central nervous system that is essential for many bodily and mental processes.^2–4^ Dopamine is involved in a delicate balance between other neurotransmitters to facilitate coordination between millions of nerves and muscle cell movement. A decrease in dopamine levels can cause Parkinson’s disease.^3,5^ Currently, L-DOPA (a dopamine precursor) is mainly used to revive Parkinson’s disease at an initial stage, but at the advanced level, with the loss of dopa decarboxylase enzyme, this drug ultimately loses its therapeutic effect.^6,7^ Therefore, researchers are exploring using dopamine-loaded nanoparticles to treat Parkinson’s disease, as direct dopamine delivery is required but cannot cross the blood-brain barrier on its own.^8,9^ Nanomaterials such as gold nanoparticles, liposomes, quantum dots, and other polymeric drug delivery systems have been used for loading and delivering Dopamine.^7–9^ However, concerns have arisen regarding the neurotoxicity triggered by these nanoparticles. Many nano-delivery methods suffer from neuroglial toxicity, undesired interactions, intra-brain accumulation, and reactive oxidative stress.^10^

DNA nanocages are an advanced nanocarrier category with excellent inherent biocompatibility and non-cytotoxicity. The Watson-Crick base pairing principle makes these structures highly programmable, allowing for precise control of shape, size, and versatile functionality.^11–14^ Recent research has demonstrated the wide use of DNA nanostructures as drug delivery systems for brain disease because of their ability to pass the blood-brain barrier (BBB).^15–20^ These properties indicate that DNA nanocages can potentially be used as drug-delivery systems for neurodegenerative diseases and be more suitable than conventional nanoscale drug carriers.^12,15–17^ DNA tetrahedral (TD) nanocages formed by self-assembling four synthetic ssDNA oligonucleotides exhibit remarkable mechanical stiffness, nontoxicity, and resistance to nuclease degradation.^21,22^ DNA tetrahedron provides enough space for small molecules as a nanocarrier and is a promising candidate in drug delivery and bioimaging platforms.^23,24^ It has been utilized as a nanocarrier for therapeutic drugs/small molecules^15^ such as doxorubicin^25^, methylene blue^26^, curcumin^27^, actinomycin D^28^, ampicillin^29^, vitamin B12 (VB12)^30^, and nucleic acid drug CpG.^31,32^ The tetrahedral DNA nanocages showed excellent cellular uptake rate among other shaped nanocages.^21,33^

Recent research has demonstrated the wide use of DNA tetrahedrons as drug-delivery vehicles to treat brain tumours and neurological disorders.^15,18,24^ DNA tetrahedrons with and without Angio-peptide-2 penetrated the BBB and were used in brain tumour imaging agents.^24^ D Y Yan et al. demonstrated that DNA nanostructure with and without BBB-penetrating peptide can cross the BBB in brain cancer treatment.^34^ They reported that ligand modification is not essential for DNA tetrahedron nanocages, it could still cross the BBB by endocytosis in both in vitro and in vivo models.^34^ W Cui et al. deliver vitamin B12-loaded DNA TD across BBB, which inhibits LRRK2 protein kinase and restores autophagy for treating Parkinson’s disease.^30^ In addition to the excellent drug delivery ability^23^, DNA TD Itself has various neurotherapeutic efficiencies, such as neuroprotective and neurotherapeutic effects^35^, neural stem cell proliferation, and neuronal differentiation ability.^36^ Many physical, chemical, and computational studies have reported Dopamine can bind to a double helical linear DNA molecule.^37–39^ Therefore, based on the above literature reports, the properties of DNA tetrahedral nanocages as blood-brain barrier crossing nanoparticles^15,34^ and their excellent drug delivery efficacy^23^ and neurotherapeutic efficiencies^15,35^, we hypothesize that DNA tetrahedron can be a viable delivery system for effective delivery of Dopamine. The Dopamine-loaded DNA tetrahedron can be potential therapeutics in treating Parkinson’s disease.^15^ As the protonated Dopamine is an active form of the neuronal dopamine transporter, we used the dopamine hydrochloride to make the TD-Dopamine complex.^40^ The protonated form of Dopamine can also enhance DNA binding through electrostatic interaction with a negatively charged phosphate backbone and intrudes into the dsDNA.

The self-assembled DNA tetrahedrons (TD) were synthesized using previously reported protocols.^21,22^ The synthesis involved mixing four complementary ss-oligonucleotides of 55 bases, T1, T2, T3, and T4, in equimolar concentration in the presence of 2 mM magnesium chloride. Thermal annealing was performed for the self-assembly of nucleotide into Tetrahedron. It is performed by following these steps: starting temperature of 95°C, decreasing the temperature by 5°C, holding for 15 minutes, and repeating the process until reaching a temperature of 4°C (**Fig 1A**). The self-assembly of these four oligos into TDs was confirmed by the electrophoretic mobility shift assay, which showed the retardation in the migration of the self-assembled TDs (T1+T2+T3+T4) compared to other lower combinations or single oligo (**Fig 1G**). The particle size distribution of the self-assembled structures was observed using dynamic light scattering (DLS), which revealed an average hydrodynamic diameter of self-assembled structures are 13.18±0.39 nm (**Fig 1B**), similar to the estimated size. Then, Atomic force microscopy (AFM) was used for topographic analysis of nanostructures. AFM imaging showed the formation of homogenously distributed self-assembled tetrahedral DNA nanostructures (**Fig 1D &E**). The histogram of particle size measurement of AFM images and subsequent Gaussian curve fit revealed that the average nanostructure size is 14.04±0.38 nm (**Fig 1F**). All the above characterization results obtained agree well with each other and confirm the formation of tetrahedral nanostructures.

**Figure 1.**
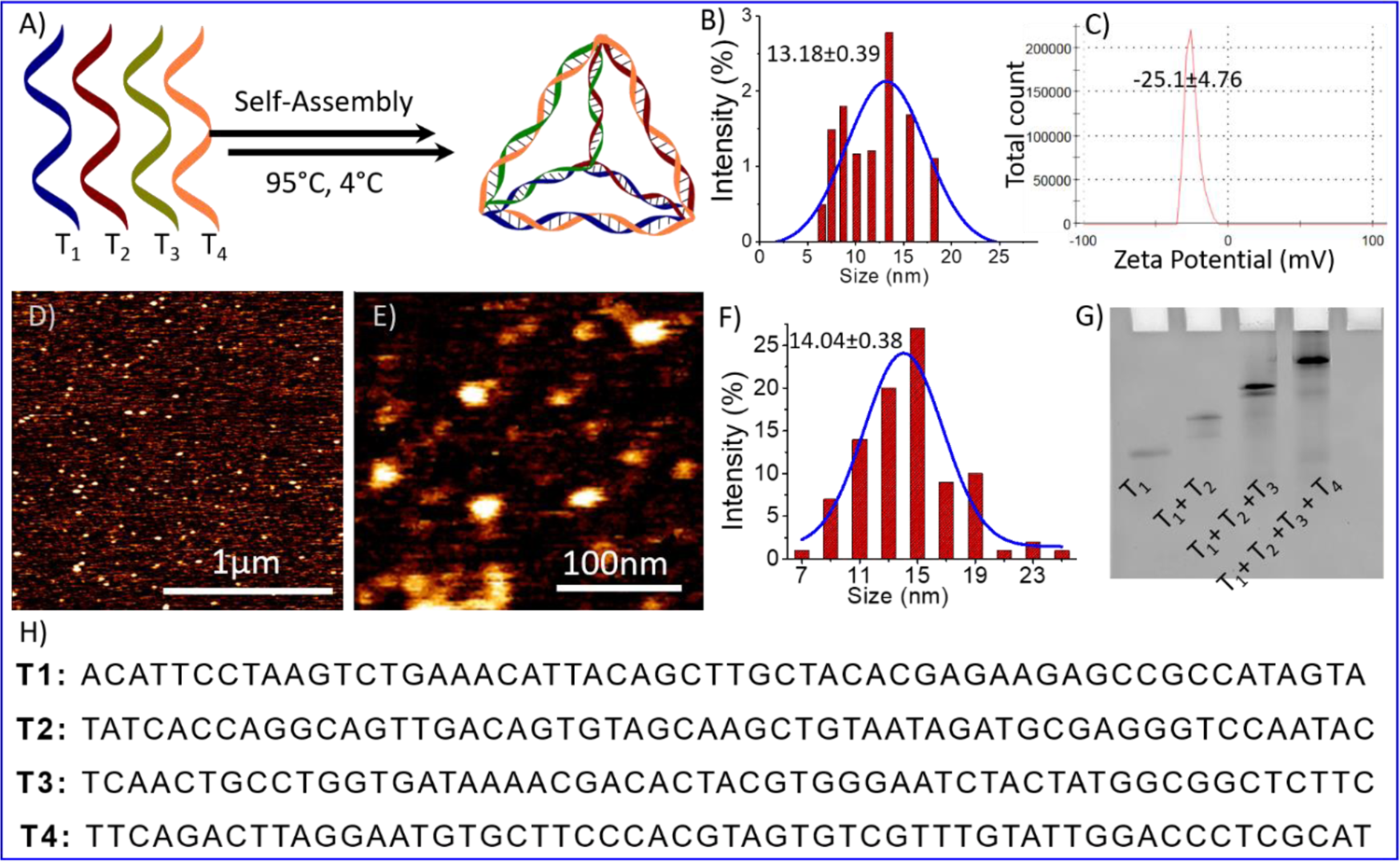
Represent formation and characterization of a tetrahedral DNA nanostructure. A) Schematic representation of the oligonucleotides self-assembly into Tetrahedron using the annealing process; B) DLS results of synthesized Tetrahedron showed the hydrodynamic diameter C) Zeta potential of negatively charged DNA nanoparticles D) AFM Image of DNA tetrahedron E) High-resolution AFM images F) bar graph showing the Particle size distribution of AFM image of TD; G) gel electrophoresis mobility shift-based characterization showing the retardation in the mobility upon formation of the Tetrahedron (T1+T2+T3+T4) and H) Oligonucleotides sequence (T1, T2, T3, and T4) used for tetrahedron synthesis.

DNA molecules can bind to various ligands or small molecules through different binding modes, including electrostatic interactions, major or minor groove space binding, and intercalation into the base pairs.^41^ Fluorescence spectroscopy is a technique used to study binding interactions without altering or destroying them.^41,42^ To investigate the binding interaction of dopamine hydrochloride with tetrahedral DNA nanostructure, a fluorescence titration experiment of dopamine hydrochloride (20µM) with the gradual addition of DNA tetrahedron (up to 80 nM) was performed. Dopamine exhibits intrinsic fluorescence emission at around 320nm on excitation at 280 nm wavelength (**Fig 2A**). Fluorescence spectra showed a quenching in fluorescence intensity on the gradual increase of DNA-TDs (**Fig 2B**). The quenching in fluorescence intensity indicates the interaction between Dopamine and TD. The Stern–Volmer constant, K_SV_ (=2.01 × 10^6^ M^-1^), has been determined from the slope of the plot of F^0^/F versus [TD] (**Fig 2I**).^41,43^ The high value of the quenching constant indicates the quenching is static due to the complex formation between DNA-TD and Dopamine.^44^

**Figure 2.**
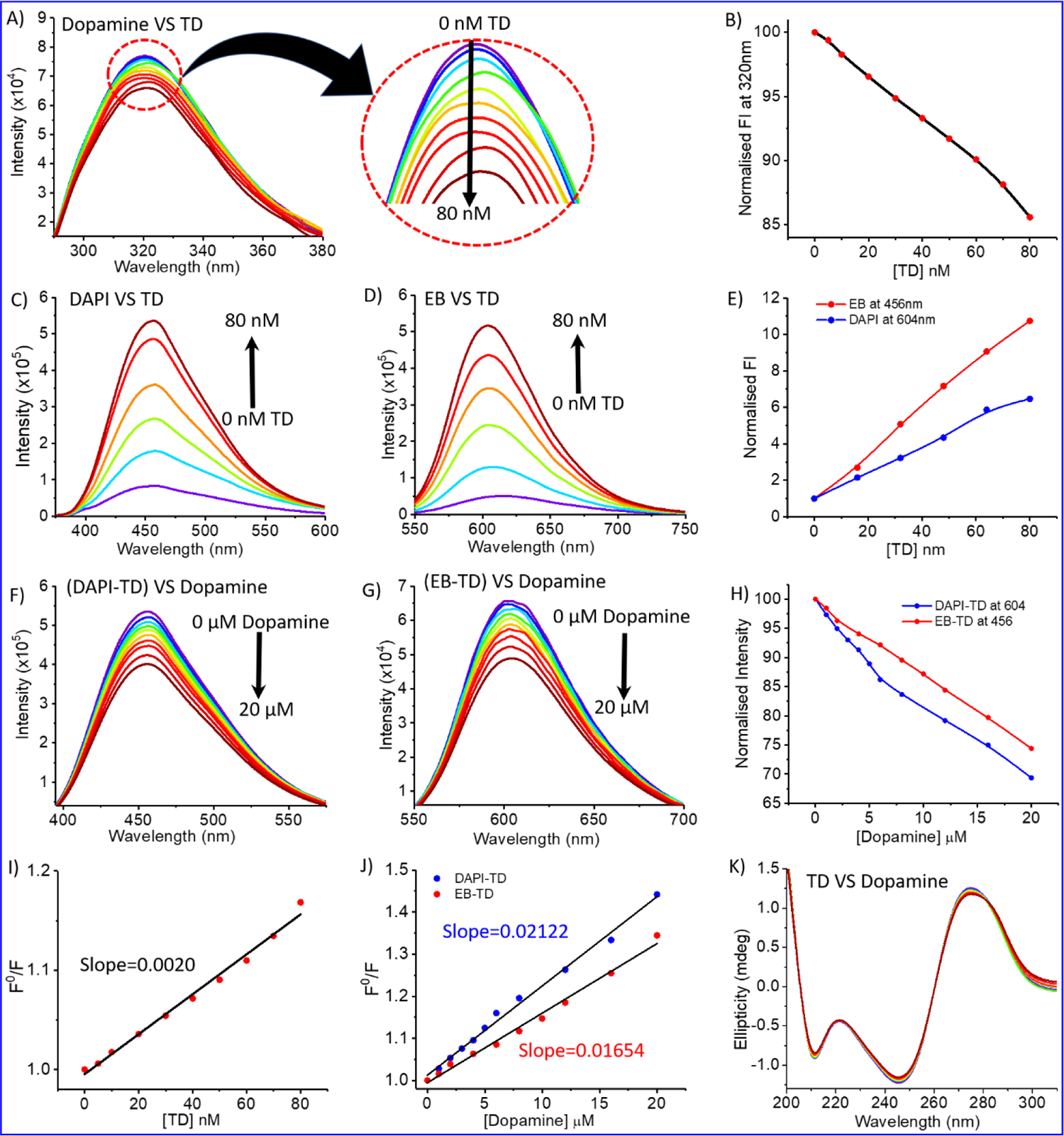
Fluorescence spectroscopy and CD analysis of DNA-Dopamine interactions. A) Fluorescence quenching spectra of Dopamine (20µM) with gradual increase of DNA TD up to 80 nM; B) Plot between fluorescence intensity of Dopamine at wavelength 320nm vs. added DNA-TD concentration up to 80 nM; C) & D) Fluorescence enhancement spectrum of DAPI (10 µM) and ethidium bromide (EB) (10 µM) on gradual increase of DNA TD up to 80nM concentration respectively. E) Plot between the fluorescence intensity of DAPI-TD complex (blue) at 456nm and EB-TD complex (red) at 604nm vs. added DNA-TD concentration up to 80 nM; F) & G) Dye displacement assay spectra of TD-DAPI and TD-EB complex by gradual addition of Dopamine respectively; H) represent the corresponding decrease in FI of TD-DAPI complex (blue) and TD-EB complex (red) on gradual addition of Dopamine; I) Stern– Volmer plots for Dopamine vs. TD titration spectra A; J) Stern–Volmer plots for dye displacement titration spectra F & G. and K) Circular Dichroism spectra of DNA TD on gradual increases of Dopamine.

Previous research proved that Dopamine binds DNA in multi-mode binding interactions via electrostatic interaction and major or minor groove binding.^39^ To get more clear information about the binding mode between dopamine chloride and tetrahedral DNA nanostructures, fluorescence quenching experiments were carried out by displacement of DNA binding dyes DAPI and ethidium bromide (EB) from DAPI-TD/EB-TD complex.^45,46^ DAPI is known to bind DNA at the groove region; however, ethidium bromide is an intercalating dye.^45,47^ Figures 2C and **2D** represent the fluoresce spectra of DAPI and ethidium bromide dyes bound with DNA tetrahedron. Only dye (DAPI/EB) has a minimum fluorescence intensity, but the addition of TD forms a complex with the dye, which increases in intensity (**Fig 2C-E**). A titration of DAPI-TD and EB-TD with the gradual addition of Dopamine decreases the fluorescence intensity of the DAPI-TD complex at 604 nm wavelength (**Fig 2F**) and the EB-TD complex at 456 nm (**Fig 2G**). This quenching of fluorescence indicates the displacement of dye (DAPI/EB) by Dopamine and the binding with DNA with their respective sites (**Fig 2H**). The displacement of DAPI suggests that Dopamine binds the minor groove side of ds-DNA edges of tetrahedron nanostructure, and displacement of EB indicates Dopamine as an intercalator. The Stern Volmer plot represents that the quenching constant for the displacement of DAPI (K=2.122 × 10^4^ M^-1^) is much greater than that of the displacement of EB (1.645 × 10^4^ M^-1^) (**Fig 2J**). That means protonated Dopamine binds the groove sides more effectively than stacked regions. The quenching constant obtained for dye displacement for DAPI and EB by Dopamine is smaller than that of Dopamine vs. TD titration. That means Dopamine also interacts with DNA TD through another mode. It may be electrostatic interaction between a negatively charged phosphate backbone and positively charged Dopamine.

Circular dichroism (CD) is a spectroscopy tool that is used to investigate conformational changes in bio-molecules such as proteins, peptides, and DNA.^48–50^ It is an absorption spectroscopy method based on the differential absorption of left and right circularly polarized light. Therefore, we used it to investigate TD’s conformational change in complexation with Dopamine. CD spectra of tetrahedral DNA nanostructures in the UV region from 190 to 300 nm showed characteristic bands of double-strand DNA, including a positive band at 275 nm and two negative bands at 245 nm and 211 nm (**Fig 2K**).^48^ The CD spectra remain almost similar on the addition of increasing concentrations of Dopamine (up to 50 µM) into the DNA Tetrahedron solution (200 nM). The lack of change in the CD spectra indicates that the complexation of Dopamine with TD does not destroy the helical confirmation or complementary base pairing. The fluorescence and CD spectral analysis suggest that Dopamine interacts with TD mainly on the groove sites.

Polyacrylamide gel electrophoresis is a widely used technique in biochemistry to analyze ligand interaction with proteins/DNA molecules. We performed an electrophoretic titration at constant Tetrahedral DNA against increasing amounts of dopamine hydrochloride to get more insight into the complexation between DNA-TD and Dopamine hydrochloride. Separate solutions of a mixture of Tetrahedron with 10 eq, 20 eq, 50 eq, 100 eq, 200 eq, and 250 eq of dopamine hydrochloride, respectively, incubated for about 30 minutes and then performed electrophoresis experiment. Ethidium bromide staining visualized that the nanostructure migration distance decreases with increasing dopamine hydrochloride concentrations in the presence of Dopamine (**Fig 3D**). Quantification of migration distance by these nanostructures to bare Tetrahedron (100%) clearly showed the retention of migration on an increasing amount of Dopamine (**Fig 3E**). In the presence of 250 equivalent of Dopamine, it is reduced to around 91% compared to TD alone. The retention in migration distance indicates an increase in the mass of the nanoparticles, which means Dopamine gets attached to the DNA tetrahedron, and a complex has been formed. Further, the DLS analysis of the TD-Dopamine complex in different ratios also proved an increase in the hydrodynamic size of tetrahedral nanostructures (**Fig S1**). The hydrodynamic size of TD in the presence of 10 eq, 20 eq, 50 eq, 100 eq, and 200 eq dopamine hydrochloride 13.21 nm, 13.88 nm, 16.5 nm, 16.91nm, 18.87 nm, 23.06nm were obtained, respectively (**Fig 3B &C**). The size increase also supports the formation of a TD-dopamine complex or successful full loading of Dopamine on TD.

**Figure 3.**
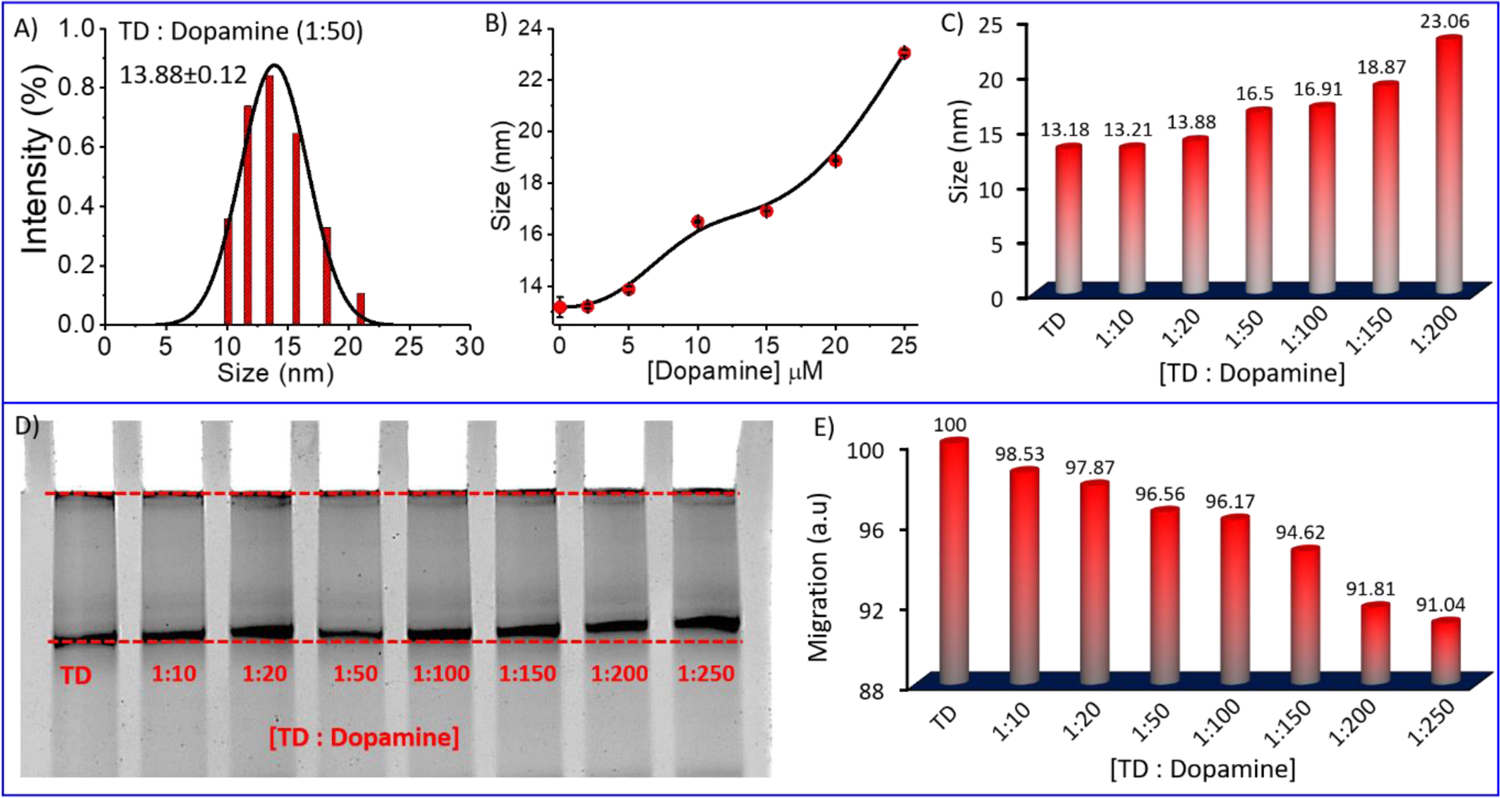
A) representative DLS result showing hydrodynamic size distribution curves for TD-Dopamine (1:50) complex; B) and C) hydrodynamic size of TD in the presence of the increasing amount of Dopamine; D) Native PAGE Electrophoretic gel-shift assay of DNA-TD in the presence of the different amount of Dopamine and E) a bar graph of corresponding electrophoretic migration of TD-Dopamine complexes in different ratio w. r. t. DNA-TD alone.

Molecular dynamics (MD) simulation is a sophisticated computational method that can be utilized to gain insights into the connections between the structure and function of macromolecules. By studying the time-dependent behaviour of microscopic systems, MD simulations can provide valuable information about the binding affinities of ligands to DNA and specific ways in which ligands bind to DNA.^51,52^ After observing the complexation of DNA and dopamine hydrochloride using biophysical and chemical techniques, we performed the MD simulations of DNA Tetrahedron for 200 ns in the presence of protonated Dopamine at physiological conditions. The initial structure of the TD was built using the polygen, and the Dopamine structure corresponding to the PubChem CID: 681, was extracted from the PubChem database.^53^ Then, the geometry optimization of the Dopamine molecule was carried out by using the B3LYP functional together with a 6-311G* basis set with the Gaussian 09 program. The built structures were then solvated in a cubic box with a buffer length of 15 Å using the TIP3P water model^54^ with xLEAP module of AMBER20.^55^ MD simulations are carried out for three systems: first TD in the presence of only one dopamine, second in the presence of 30 dopamine and 100 dopamine molecules (**Fig 4A, F & Fig S2**). The simulation of TD with single dopamine reveals that the dopamine molecule binds to the minor groove site of DNA-TD and remains there throughout the 200 ns long simulation. Our simulations with higher number of dopamine (30 and 100) suggest that Dopamine can bind both at the minor and major groove of TD and also at the phosphate backbone of DNA-TD (**Fig 4B**). The histogram of the minimum distance of Dopamine from DNA-TD shows that 28 out of 30 dopamine molecules bind the Tetrahedron (**Fig 4E**). Further, the radial distribution function (RDF) of the hydroxyl group and amine group of Dopamine and the DNA-TD suggests that protonated Dopamine interacts with minor grooves through hydroxyl and NH3+ groups via hydrogen bonding; in major grooves, it binds mainly through NH_3_^+^ groups (**Fig 4C &D**). In the case of 100 dopamine molecules, we also found the multi-mode binding of dopamine molecules over the DNA tetrahedron. The histogram of the minimum distance of Dopamine from tetrahedral nanostructures shows that 82 molecules bond to the DNA tetrahedron, and 18 are far from it (**Fig 4F & G**).

**Figure 4.**
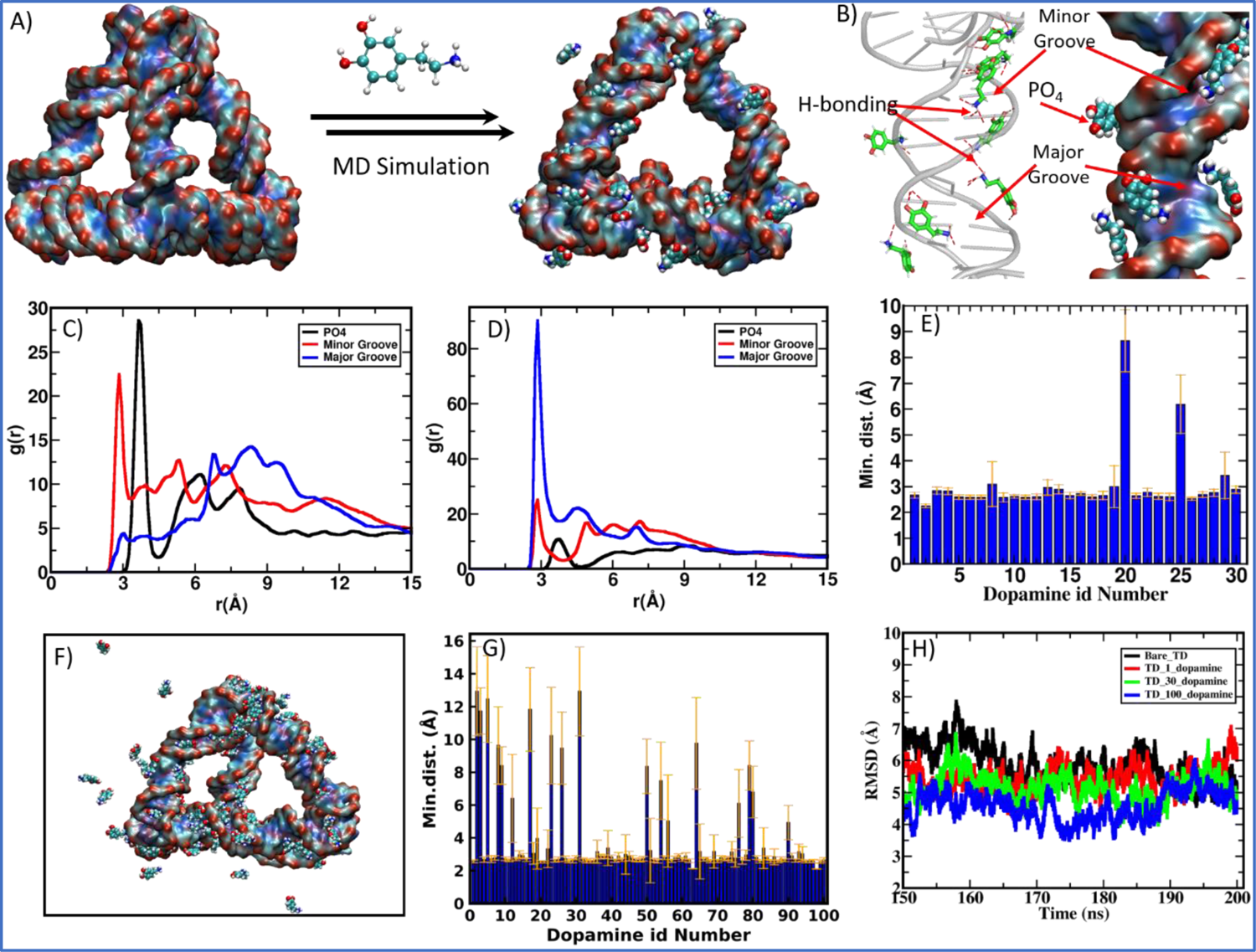
Molecular Dynamics Simulation (MD) study of Dopamine and DNA Tetrahedron nanocages complexation. A) Instantaneous snapshot of bare DNA-TD nanocage and TD-Dopamine complex after 200 ns long MD simulation of TD with 30 protonated dopamine molecules; B) represents high resolution image of dopamine binding interactions to the DNA-TD at minor groove, major groove and with phosphate (PO_4_) back bone from the MD simulation configuration C) Radial distribution functions (RDF) between the oxygen atoms of the hydroxyl group of Dopamine and the P atom of phosphate, N6, O4, and N4 electronegative atoms of bases for the Major Groove and N3 and O2 electronegative atoms of bases for the Minor Groove of TD; D) RDF between the N atoms of the amine group of Dopamine and the P atom of phosphate, N6, O4, and N4 electronegative atoms of bases for the Major Groove; and N3 and O2 electronegative atoms of bases for the Minor Groove of TD; E) Histogram for the minimum distance between DNA-TD and 30 protonated dopamine molecules after 200ns simulation; F) TD-Dopamine complex after MD simulation for 200 ns of TD with 100 protonated dopamine molecules; G) Histogram for the minimum distance between DNA-TD and 100 protonated dopamine molecules after 200ns simulation, and H) Root-mean-square deviation (RMSD) during last 50ns simulation of TD with and without Dopamine.

The instantaneous snapshot of the TD-Dopamine mixture at the end of 200 ns long MD simulation showed that the helical structure of ds-DNA is not affected by binding the dopamine molecules, as also confirmed by our spectroscopic observation. For closer insight into the effect of Dopamine on TD structure, we calculate the root mean squire deviation (RMSD) for the entire simulation period. RMSD as a function of simulation time showed that in the presence of 100 protonated dopamine molecules, RMSD is slightly lower than that of bare TD alone (**Fig 4H**). So, in the presence of dopamine molecule, the DNA TD structure is slightly more stable compared to bare TD. Otherwise, DNA-TD structure maintain its canonical form both in the absence and presence of dopamine molecules. We also calculate the degree of hydrogen bonding (base pairing) responsible for the self-assembly of oligos into the tetrahedral nanostructure. The number of hydrogen bonds for bare TD is 246.8±1.2, while in the presence of 100 dopamine, it increases to 253.7±2.1. The increase in hydrogen bonding further supports the stabilization of DNA tetrahedron in the presence of dopamine molecules.

Considering the analysis mentioned above, we investigate TD’s morphological change in the presence of Dopamine using atomic force microscopy (AFM). We added dopamine hydrochloride in the solution of DNA-TD in different ratios separately and incubated them for 2h before AFM imaging. AFM imaging of the TD-dopamine complex revealed that the size of nanostructures increases from 13 nm to 23 nm with increasing Dopamine concentration (Fig 5). The AFM image of DNA tetrahedral nanostructures in the presence of 20 eq of Dopamine showed a slight increase in size; however, no significant change was observed in morphology (**Fig 5A**). A particle size distribution was done with the help of images followed by Gaussian fitting in origin pro, revealing the size range from 7 to 29 nm with an average size of 15.66±0.83 nm (**Fig 5B**). In the presence of 50eq, Dopamine also showed similar nanoparticles of the size ranging from 8-38 nm with an average diameter of 18.21±0.55nm (**Fig 5C &D**). The morphology of the TD-Dopamine complex, similar to DNA-TD nanostructures, suggests that most of the dopamine molecules bind with the ds-DNA edge of the Tetrahedron without disturbing the original conformation of TD. These AFM results have good agreement results obtained from spectroscopy and simulation analysis. With further increases in dopamine concentrations, the morphology of TD-dopamine seems to transform into spherical or distorted sphere-like structures (**Fig 5E**). The average particle size of the TD-dopamine complex (1:100) was around 19.92±0.71 nm (**Fig 5F**). In the case of the TD-dopamine complex in a 1:200 ratio, the atomic force microscopy image revealed the formation of homogeneously distributed sphere-like structures (**Fig 5G**). The particle size distribution of the AFM image revealed an increase in the average size of the nanostructures, which is 23.17±0.82nm (**Fig 5H**). Our MD simulation study showed that few dopmaine molecules remain unbound in the case of 100 dopamine molecules during the 200 ns long dynamics. More prolonged incubation of Dopamine with DNA-TD may allow them to deposit onto the negative backbone of TD. The TD nanoparticles get coated with dopamine molecules on increased Dopamine concentration, which is also shown in the AFM images.

**Figure 5.**
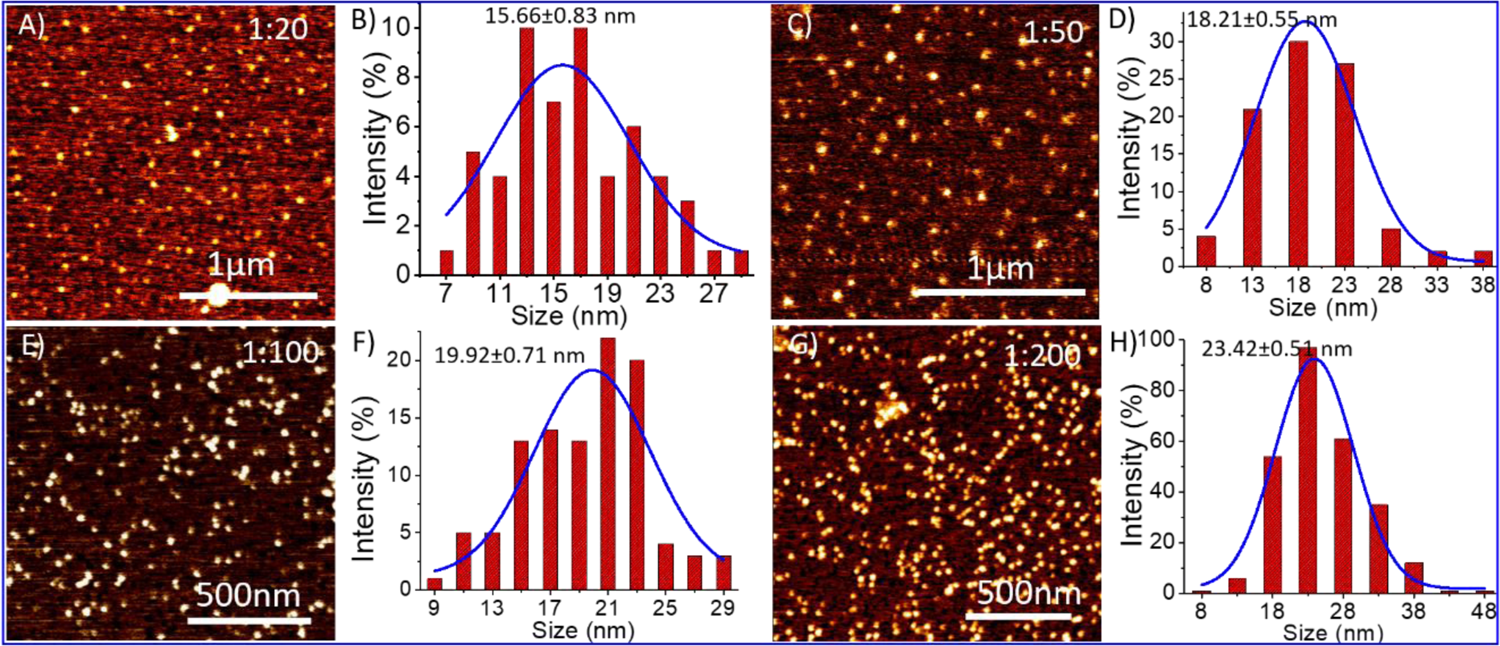
Atomic force microscopy (AFM) results of TD and TD-Dopamine complexes nanostructures at different concentrations of Dopamine, and their particle size analysis A) AFM image of TD-Dopamine (1:20); B) Corresponding bar graph of size distribution analysis; C) AFM image of TD-Dopamine (1:50); D) Corresponding bar graph of size distribution analysis; E) AFM image of TD-Dopamine (1:100); F) Corresponding bar graph of size distribution analysis; G) AFM image of TD-Dopamine (1:200); and H) Corresponding bar graph of size distribution analysis.

The study of DNA TD-dopamine complexation from spectroscopy, microscopy, and computational simulation has proved that DNA-Tetrahedron can load a high amount of Dopamine and be used as a dopamine carrier. DNA and Dopamine are crucial components in the human body and possess inherent biocompatibility. However, a cell viability experiment using the MTT assay was conducted to assess the cytotoxicity of TD-Dopamine complex nanoparticles on SH-SY5Y cells, and the results showed no significant decrease in cellular viability (**Fig 6D &E**). Human SH-SY5Y cells have been extensively used to study in-vitro neurodegenerative illnesses and as an in vitro model for neuroscience and neurotoxicity investigations.^56,57^ The SH-SY5Y human neuroblastoma cell line, either undifferentiated or developed into neuron-like cells, is one of the most often used cell lines in neuroscience.^56,57^ Cellular internalization of the nanostructure is a primary step to work as a delivery system; therefore, we did the cellular uptake study of TD-dopamine nanostructures on both differentiated and undifferentiated SH-SY5Y neuroblastoma cells. In the cellular uptake study, one of the four oligonucleotides (T4) was labeled with cyanine-3 (cy3) dye at 5′ ends for tracking purposes. Our previous report found that Cy3-label DNA tetrahedron (150 nM) showed significant uptake in SH-SY5Y cells. Therefore, we used 150nM DNA Tetrahedron alone and with different amounts of dopamine hydrochloride, 50eq, 100eq, 150eq, and 200eq, for cell treatment (**Fig 6A & B**). The SHSY5Y cells were incubated with all these combinations for 15 min at 37°C and subjected the samples to laser scanning confocal microscopy after nucleus staining with Hoechst. Negligible signals were observed in the red channel for untreated cells and cells treated with Dopamine (30µM, 200eq of [TD]) on confocal imaging (**Fig 6A**). The confocal image of the cells treated with TD and TD-dopamine complexes clearly showed the presence of Cy3 labeled TD. A quantification analysis of Cy3 signal intensity of confocal images reveals that the Dopamine-loaded DNA-TD nanostructures are significantly uptaken by cells. In the case of 1:100 (TD-dopamine) combinationtion, maximum cellular uptake is found (**Fig 6C**). Similar trends were found in the cellular uptake study in nondifferentiated SHSY5Y cells (**Fig S3**).

**Figure 6.**
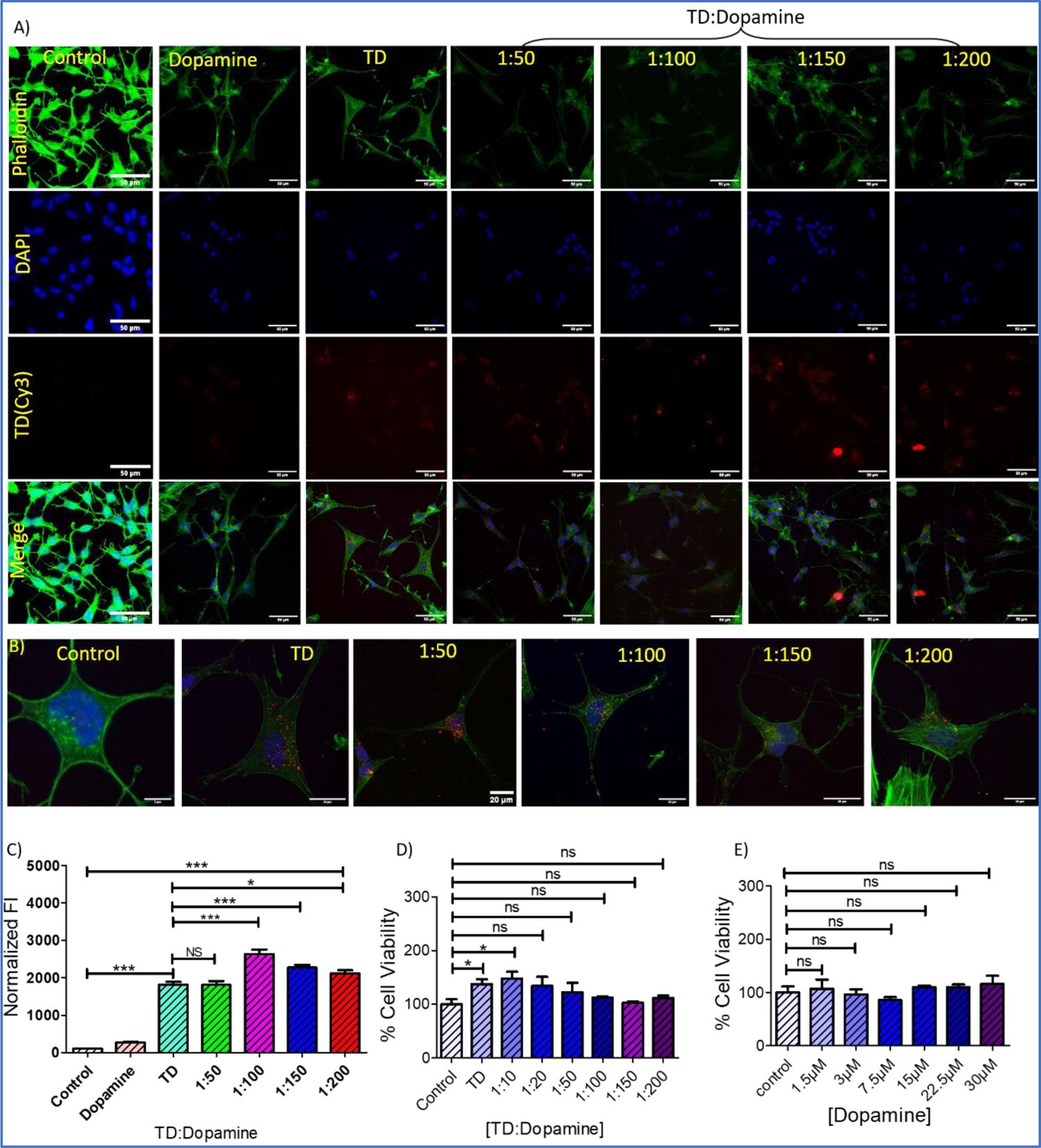
Cellular uptake of TD-Dopamine complex nanostructures into differentiated SH-SY5Y neuroblastoma cell lines. A) Confocal imaging of differentiated SH-SY5Y neuroblastoma cells treated with DNA Cy3 TD (150 nm) and with Cy3 TD (150 nm)– Dopamine in different ratios of TD and Dopamine for 15 min. The green channel represents an actin cytoskeleton stained with Phalloidin-A488, the blue channel represents nuclei stained with DAPI, the red channel represents TD(Cy3) uptake, and the lower panel represents merged images. The scale bar is 50μm. (B) Magnified confocal images of the treated and untreated cells. The scale bar is 20 μm; C) Quantifying TD–Cy3 uptake in differentiated SHSY-5Y neuroblastoma cells from panel A. ***Statistically significant p-value (p < 0.0001) and *Statistically significant p-value (P<0.05). D) and E) Bar graphs representing cell viability (%) obtained from the MTT assay test against different concentrations of Dopamine with TD (150nm) and without TD. *Statistically significant p-value (P<0.05).

The ability to differentiate SH-SY5Y neuroblastoma cells into a more mature, neuron-like phenotype through manipulating the culture medium has provided numerous benefits in neuroscience research.^56,57^ A progressive death of dopaminergic neurons (dopamine-synthesizing neurons) causes a reduction of dopamine levels in the brain striatum. The differentiation of dopaminergic cells can boost the treatment of Parkinson’s disease.^58,59^ Therefore, cell differentiation also plays an essential role in treating neurological diseases. DNA tetrahedral nanostructures, known as neuronal cell differentiation, for example, tetrahedral nanostructures reported to promote NE-4C stem cell differentiation and proliferation.^36^ Therefore, an experiment was performed with DNA-TD, Dopamine, and different combinations of TD-dopamine complex to check the ability to differentiate SH-SY5Y cells. The differentiation experiment was done by following a well-established protocol.^56,60,61^ Around 10^5^ cells per well were seeded on a 10mm glass coverslip in a four-well plate 24h before the experiment. On the first day, cells were treated with DNA-TD, dopamine TD-dopamine complex at different differentiation media (DM)-1 composition. Untreated cells cultured only in DM-1 were used as a control, and retinoic acid was used as a standard differentiator. The same treatment was repeated on day 3 and day 5. On day 7 and day 9, the same treatment was done in DM-2, and on day 11 was treated in DM-3.

The growth and morphological changes due to differentiation were regularly monitored using an optical microscope (**Fig S4-S9**). The confocal image showed morphological differences between undifferentiated and differentiated SH-SY5Y cells (**Fig 7A**). The undifferentiated cells have fewer neuritic projections with a linear type morphology; however, the differentiated cells showed multiple neuritic arms with larger area inflated morphology. The confocal images revealed that DNA-TD alone and combined with Dopamine can differentiate the SH-SY5Y cells. The treatment with free Dopamine also showed differentiation of SH-SY5Y cells. Quantification of these differentiated cells showed that retinoic acid, a standard differentiating agent, differentiates all the cells into neurons (**Fig 7B**). However, the cells treated with TD, Dopamine, and TD-dopamine combinations also showed significant differentiation compared to control. Optical microscopy images also found some of the differentiated cells are like typical neurons (**Fig 7C**). These cellular studies proved that DNA-TD can serve as a nanocarrier as well as a differentiating agent. These results, combined with the efficiency of TD nanostructures to cross the blood-brain barrier, suggest that dopamine-loaded DNA-TD can be an excellent nano-delivery system for Parkinson’s disease treatment. This approach can potentially improve Dopamine delivery to the brain and enhance its therapeutic efficacy.

**Figure 7.**
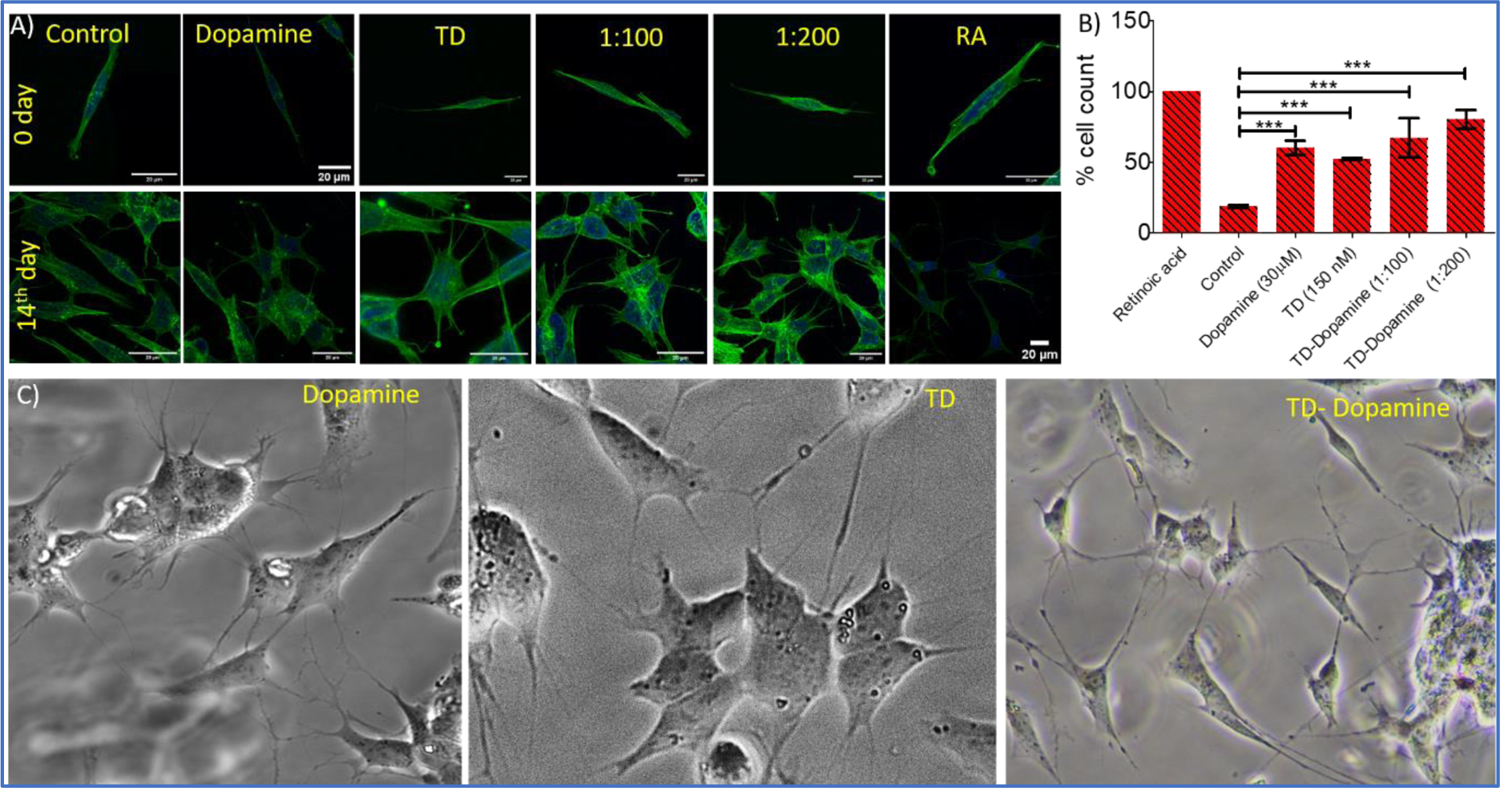
Confocal imaging of SHSY5Y cells differentiation study. A) The upper panel represents undifferentiated SHSY5Y cells, and the lower panel represents differentiated SH-SY5Y cells on the 14^th^ day. B) quantification of % cell count differentiated concerning total cell number and C) Representative optical microscopy images of differentiated cells. ***Statistically significant p-value (p < 0.0001).

Zebrafish larvae are a commonly used model organism in research due to their transparency, high fecundity, and suitable brain dimensions. They are used in various studies, such as toxicity testing, behavioural assays, circuit modelling, and uptake studies.^62,63^ The zebrafish genome shares 70% of our genetic makeup, making it a superior *in vivo* model for studying how different biological substances are absorbed.^64^ Our previous study showed that TD did not cause any alterations in the survival rate and hatching success of developing zebrafish embryos.^65^ Study also demonstrated a significant uptake of DNA-TD compared to other geometries of DNA nanocages.^65,66^ At this time, to assess the biocompatible nature of TD and TD-Dopamine complex on developing zebrafish larvae, the 8 dpf zebrafish larvae were exposed to TD (300 nM) and TD-Dopamine (1:100). The survival rate, malformation, and heart rate were assessed for the period of 6 and 24 hours, respectively. We observed that TD and TD-Dopamine exhibited a biocompatible nature toward the developing zebrafish larvae post 6 hours of treatments (**Fig 8C**). The survival rate for TD and TD-Dopamine from zero to 6 hours was the same as for controls. Dopamine also exhibited a biocompatible nature towards developing zebrafish larvae until 6 hours of exposure. No malformation was observed in larvae until 24 hours of exposure to TD, Dopamine, and TD-dopamine (1:100) (**Fig 8D**). The heart rate per minute of developing larvae was 150, 142, 154, and 138 for control, TD, Dopamine, and TD-Dopamine (**Fig 8E**). However, the initial heart rates in larvae exposed by alone Dopamine were 175/minute till 3 hours of exposure, and the highly active behaviour of larvae was also observed. The results depict that the TD and TD-Dopamine exerted no toxic effect on the survival and heart rate of the zebrafish larvae compared to the control.

**Figure 8.**
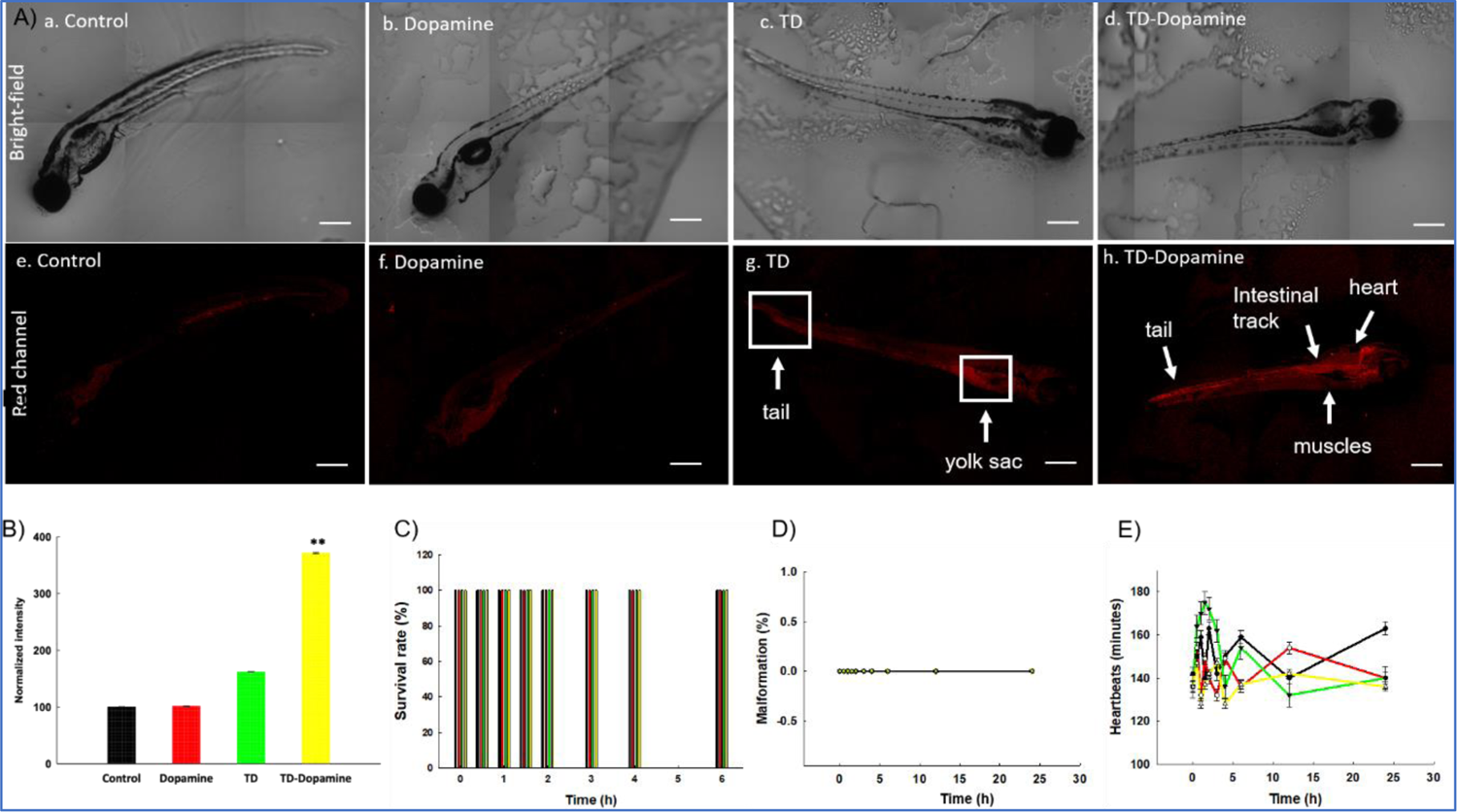
Uptake analysis of TD, Dopamine, and TD-dopamine in zebrafish embryos. (A) Uptake of TD, Dopamine, and TD-dopamine (1:100) in zebrafish embryos post 4 h of treatment. (a−d) Brightfield images of zebrafish embryos post 4 h of treatment. (e−h) Red channel images of zebrafish embryos post 4 h of treatment. (B) Fluorescence intensity and quantification analysis of TD, Dopamine, and TD−Dopamine uptake in zebrafish embryos post 4 h of treatment. The scale bar is set at 50 μm. **Statistically significant p-value (p < 0.01). Ten embryos per condition were quantified. Data represent the mean SEM of three independent experiments. Effects of TD (300 nM), Dopamine, and TD−Dopamine (1:100) on the survival rate C), malformations D) and heartbeats E) of zebrafish embryos from 4 hours to 24 hours post treatments. Data represent the mean SEM of three independent experiments.

The uptake capacity of any nanocarrier is crucial for its potential biomedical applications. Further, we proceed to analyze and quantify the uptake potential of TD-Dopamine in zebrafish larvae. The larvae were exposed to TD (300nM) and TD-Dopamine complex (1:100) for 4 hours of treatment (**Fig 8A**). TD and TD-Dopamine (1:100) uptake was quantified using a laser-scanning confocal microscope. One of the four oligonucleotides (T4) was labelled with cyanine-5 (Cy5) dye at their 5′ ends for tracking purposes. The signal for TD-Dopamine (1:100) was significantly higher than alone TD in larvae post 4 h treatment (**Fig 8A**). Additionally, the precise location of the targeted organelles *in vivo* depends on the physiological mechanism of intracellular transport and the unique characteristics of the subcellular structures. The usefulness of DNA nanostructures for imaging organelles for diagnostic applications has been demonstrated by precise targeting at the organelle level using fluorescently tagged DNA nanodevices; the same can be examined for therapeutic applications. Understanding the rules of appropriate transmission between organelles, cells, and tissues may help develop DNA nanodevices as diagnostic and therapeutic tools. Moreover, our uptake studies indicated that dopamine-modified TD enhanced the uptake and internalization in zebrafish larvae. The confocal image of larvae clearly shows the uptake and internalization of the TD-Dopamine nanostructures. Quantification analysis of Cy5 signal intensity of TD-Dopamine exerted the significant uptake with 100 eq. Dopamine compared to TD alone (**Fig 8B**).

## Conclusion

In this work, we studied the complexation mechanism of DNA tetrahedron nanocage and dopamine hydrochloride. The results showed that Dopamine binds with the tetrahedral nanostructure through hydrogen bonding and electrostatic interaction. Through fluorescence spectroscopy analysis, we confirmed the binding of Dopamine with the tetrahedral nanostructure. The dye displacement experiments and CD spectral observations showed that Dopamine primarily binds to the groove region of DNA-TD without disrupting the Tetrahedron’s original helical structures of edges. Molecular dynamic simulations further supported the binding of protonated Dopamine in both the major and minor groove regions, along with the negatively charged phosphate backbone. A combination of hydrogen bonding and electrostatic interactions facilitated this binding. Simulation results indicated that out of 100 dopamine molecules, approximately 82 molecules bound to TD within a 200ns simulation. The binding of Dopamine increased the degree of hydrogen bonding between DNA base pairs, thereby enhancing the stability of TD. Further, the dynamic light scattering analysis reveals an increase in the hydrodynamic diameter of nanostructures on increasing Dopamine. Electrophoretic migration gel analysis and AFM imagining further supported the increased size. The rise in nanostructure size may be the deposition of dopamine molecules on the DNA backbone of TD through electrostatic interactions. They were subjected to biological application after successfully loading Dopamine onto the TD. The Dopamine-loaded TD nanostructures demonstrated significant cellular internalization with differentiated and nondifferentiated SH-SY5Y neuroblastoma cells. Further, it was found that DNA-TD and TD-Dopamine complexes promote the differentiation of SH-SY5Y neuroblastoma cells. Additionally, we demonstrated the bio-compatibility and uptake of TD-Dopamine complex *in vivo* model zebrafish larvae. The result showed a significant uptake of TD and TD-dopamine complex in zebrafish larvae, and no significant changes were obtained regarding survival rate, malformations, and heartbeats. These findings imply that dopamine-loaded DNA-TD can be a highly effective nano-delivery system for treating Parkinson’s disease, along with high biocompatibility, non-cytotoxicity, and the effectiveness of TD nanostructures in crossing the blood-brain barrier. This strategy may enhance Dopamine’s therapeutic efficacy by improving its transport to the brain.

## Acknowledgments

RS thanks, Gujcost-DST and IITGN-MoE, GoI, for the Postdoctoral Fellowship. KK thanks SERB, and GoI, for the National Postdoctoral Fellowship. SM thanks CSIR, India for research fellowship. PKM thanks SERB, India for funding. DB thanks SERB, and GoI for the Ramanujan Fellowship, IITGN for the start-up grant, and DBT-EMR, Gujcost-DST, and GSBTM for research grants. We thank CIF ITTGN for the instrumental facilities.

## References

1 S. E. Hyman, Current Biology, 2005, 15, R154–R158.

2 R. I. Teleanu, A.-G. Niculescu, E. Roza, O. Vladâcenco, A. M. Grumezescu and D. M. Teleanu, Int J Mol Sci, 2022, 23, 5954.

3 H. Juárez Olguín, D. Calderón Guzmán, E. Hernández García and G. Barragán Mejía, Oxid Med Cell Longev, 2016, 2016, 1–13.

4 D. L. Robinson, A. Hermans, A. T. Seipel and R. M. Wightman, Chem Rev, 2008, 108, 2554– 2584.

5 F. N. Emamzadeh and A. Surguchov, Front Neurosci, DOI:10.3389/fnins.2018.00612.

6 D. Salat and E. Tolosa, J Parkinsons Dis, 2013, 3, 255–269.

7 R. Pahuja, K. Seth, A. Shukla, R. K. Shukla, P. Bhatnagar, L. K. S. Chauhan, P. N. Saxena, J. Arun, B. P. Chaudhari, D. K. Patel, S. P. Singh, R. Shukla, V. K. Khanna, P. Kumar, R. K. Chaturvedi and K. C. Gupta, ACS Nano, 2015, 9, 4850–4871.

8 J. Baskin, J. E. Jeon and S. J. G. Lewis, J Neurol, 2021, 268, 1981–1994.

9 V. Monge-Fuentes, A. Biolchi Mayer, M. R. Lima, L. R. Geraldes, L. N. Zanotto, K. G. Moreira, O. P. Martins, H. L. Piva, M. S. S. Felipe, A. C. Amaral, A. L. Bocca, A. C. Tedesco and M. R. Mortari, Sci Rep, 2021, 11, 15185.

10 D. Teleanu, C. Chircov, A. Grumezescu and R. Teleanu, Nanomaterials, 2019, 9, 96.

11 W. Ma, Y. Zhan, Y. Zhang, C. Mao, X. Xie and Y. Lin, Signal Transduct Target Ther, 2021, 6, 351.

12 Q. Hu, H. Li, L. Wang, H. Gu and C. Fan, Chem Rev, 2019, 119, 6459–6506.

13 A. R. Gada, P. Vaswani, R. Singh and D. Bhatia, ChemBioChem,, DOI:10.1002/cbic.202200634.

14 D. Menon, R. Singh, K. B. Joshi, S. Gupta and D. Bhatia, ChemBioChem,, DOI:10.1002/cbic.202200580.

15 X. Chen, Y. Xie, Z. Liu and Y. Lin, Front Bioeng Biotechnol,, DOI:10.3389/fbioe.2021.782237.

16 X. Chen, D. Liu, Y. Wu, H. Yao, Q. Xia and Y. Yang, ChemBioChem,, DOI:10.1002/cbic.202200459.

17 P. Hivare, C. Panda, S. Gupta and D. Bhatia, ACS Chem Neurosci, 2021, 12, 363–377.

18 J. Yan, X. Zhan, Z. Zhang, K. Chen, M. Wang, Y. Sun, B. He and Y. Liang, J Nanobiotechnology, 2021, 19, 412.

19 F. Xiao, L. Lin, Z. Chao, C. Shao, Z. Chen, Z. Wei, J. Lu, Y. Huang, L. Li, Q. Liu, Y. Liang and L. Tian, Angewandte Chemie International Edition, 2020, 59, 9702–9710.

20 S. Shi, W. Fu, S. Lin, T. Tian, S. Li, X. Shao, Y. Zhang, T. Zhang, Z. Tang, Y. Zhou, Y. Lin and X. Cai, Nanomedicine, 2019, 21, 102061.

21 A. Rajwar, S. R. Shetty, P. Vaswani, V. Morya, A. Barai, S. Sen, M. Sonawane and D. Bhatia, ACS Nano, 2022, 16, 10496–10508.

22 N. M. Rao, Chem Phys Lipids, 2010, 163, 245–252.

23 Y. Hu, Z. Chen, H. Zhang, M. Li, Z. Hou, X. Luo and X. Xue, Drug Deliv, 2017, 24, 1295–1301.

24 T. Tian, J. Li, C. Xie, Y. Sun, H. Lei, X. Liu, J. Xia, J. Shi, L. Wang, W. Lu and C. Fan, ACS Appl Mater Interfaces, 2018, 10, 3414–3420.

25 K.-R. Kim, D.-R. Kim, T. Lee, J. Y. Yhee, B.-S. Kim, I. C. Kwon and D.-R. Ahn, Chemical Communications, 2013, 49, 2010.

26 N. Chen, S. Qin, X. Yang, Q. Wang, J. Huang and K. Wang, ACS Appl Mater Interfaces, 2016, 8, 26552–26558.

27 M. Zhang, X. Zhang, T. Tian, Q. Zhang, Y. Wen, J. Zhu, D. Xiao, W. Cui and Y. Lin, Bioact Mater, 2022, 8, 368–380.

28 M. I. Setyawati, R. V. Kutty, C. Y. Tay, X. Yuan, J. Xie and D. T. Leong, ACS Appl Mater Interfaces, 2014, 6, 21822–21831.

29 Y. Sun, S. Li, Y. Zhang, Q. Li, X. Xie, D. Zhao, T. Tian, S. Shi, L. Meng and Y. Lin, ACS Appl Mater Interfaces, 2020, 12, 36957–36966.

30 W. Cui, X. Yang, X. Chen, D. Xiao, J. Zhu, M. Zhang, X. Qin, X. Ma and Y. Lin, Adv Funct Mater,, DOI:10.1002/adfm.202105152.

31 H. Lee, A. K. R. Lytton-Jean, Y. Chen, K. T. Love, A. I. Park, E. D. Karagiannis, A. Sehgal, W. Querbes, C. S. Zurenko, M. Jayaraman, C. G. Peng, K. Charisse, A. Borodovsky, M. Manoharan, J. S. Donahoe, J. Truelove, M. Nahrendorf, R. Langer and D. G. Anderson, Nat Nanotechnol, 2012, 7, 389–393.

32 J. Li, H. Pei, B. Zhu, L. Liang, M. Wei, Y. He, N. Chen, D. Li, Q. Huang and C. Fan, ACS Nano, 2011, 5, 8783–8789.

33 R. Duangrat, A. Udomprasert and T. Kangsamaksin, Cancer Sci, 2020, 111, 3164–3173.

34 D. Y. Tam, J. W.-T. Ho, M. S. Chan, C. H. Lau, T. J. H. Chang, H. M. Leung, L. S. Liu, F. Wang, L. L. H. Chan, C. Tin and P. K. Lo, ACS Appl Mater Interfaces, 2020, acsami.0c02957.

35 W. Cui, Y. Zhan, X. Shao, W. Fu, D. Xiao, J. Zhu, X. Qin, T. Zhang, M. Zhang, Y. Zhou and Y. Lin, ACS Appl Mater Interfaces, 2019, 11, 32787–32797.

36 W. Ma, X. Shao, D. Zhao, Q. Li, M. Liu, T. Zhou, X. Xie, C. Mao, Y. Zhang and Y. Lin, ACS Appl Mater Interfaces, 2018, 10, 7892–7900.

37 K. Skúpa, M. Melicherčík and J. Urban, J Mol Model, 2015, 21, 241.

38 J. Liu, Z.-H. Wang, G.-A. Luo, Q.-W. Li and H.-W. Sun, Analytical Sciences, 2002, 18, 751–755.

39 S. Sarkar, A. Chowdhury and P. C. Singh, J Phys Chem B, 2019, 123, 10700–10708.

40 J. L. Berfield, L. C. Wang and M. E. A. Reith, Journal of Biological Chemistry, 1999, 274, 4876– 4882.

41 M. Sirajuddin, S. Ali and A. Badshah, J Photochem Photobiol B, 2013, 124, 1–19.

42 M. Khorasani-Motlagh, M. Noroozifar, A. Moodi and S. Niroomand, J Photochem Photobiol B, 2013, 127, 192–201.

43 R. Singh, N. K. Mishra, P. Gupta and K. B. Joshi, Chem Asian J, 2020, 15, 531–539.

44 R. Singh, N. Kumar Mishra, V. Kumar, V. Vinayak and K. Ballabh Joshi, ChemBioChem, 2018, 19, 1630–1637.

45 L. A. Reis and M. S. Rocha, Biopolymers, 2017, 107, e23015.

46 E. F. Healy, J Chem Educ, 2007, 84, 1304.

47 P. O. Vardevanyan, A. P. Antonyan, M. A. Parsadanyan, H. G. Davtyan and A. T. Karapetyan, Exp Mol Med, 2003, 35, 527–533.

48 V. I. Dodero, Frontiers in Bioscience, 2011, 16, 61.

49 V. Kumar, R. Singh and K. B. Joshi, New Journal of Chemistry, 2018, 42, 3452–3458.

50 R. Singh, V. Suryavashi, V. Vinayak and K. B. Joshi, ChemistrySelect, 2019, 4, 5810–5816.

51 V. Maingi, M. V. S. Kumar and P. K. Maiti, J Phys Chem B, 2012, 116, 4370–4376.

52 A. K. Sahoo, B. Bagchi and P. K. Maiti, J Chem Phys, 151, 164902 (2019).

53 C. Alves, F. Iacovelli, M. Falconi, F. Cardamone, B. Morozzo della Rocca, C. L. P. de Oliveira and A. Desideri, J Chem Inf Model, 2016, 56, 941–949.

54 W. L. Jorgensen, J. Chandrasekhar, J. D. Madura, R. W. Impey and M. L. Klein, J Chem Phys, 1983, 79, 926–935.

55 Case, David A., et al. “Amber 2020: University of california.” San Francisco (2020).

56 J. Kovalevich and D. Langford, 2013, pp. 9–21.

57 H. Xicoy, B. Wieringa and G. J. M. Martens, Mol Neurodegener, 2017, 12, 10.

58 F. N. Emamzadeh and A. Surguchov, Front Neurosci,, DOI:10.3389/fnins.2018.00612.

59 P. P. Michel, E. C. Hirsch and S. Hunot, Neuron, 2016, 90, 675–691.

60 M. M. Shipley, C. A. Mangold and M. L. Szpara, Journal of Visualized Experiments,, DOI:10.3791/53193.

61 P. Hivare, A. Gangrade, G. Swarup, K. Bhavsar, A. Singh, R. Gupta, P. Thareja, S. Gupta and D. Bhatia, Nanoscale, 2022, 14, 8611–8620.

62 S. Cassar, I. Adatto, J. L. Freeman, J. T. Gamse, I. Iturria, C. Lawrence, A. Muriana, R. T. Peterson, S. Van Cruchten and L. I. Zon, Chem Res Toxicol, 2020, 33, 95–118.

63 R. Basnet, D. Zizioli, S. Taweedet, D. Finazzi and M. Memo, Biomedicines, 2019, 7, 23.

64 K. Mhalhel, M. Sicari, L. Pansera, J. Chen, M. Levanti, N. Diotel, S. Rastegar, A. Germanà and G. Montalbano, Cells, 2023, 12, 252.

65 K. Kansara, R. Singh, P. Yadav, A. Mansuri, A. Kumar and D. Bhatia, ACS Appl Nano Mater, 2023, 6, 13443–13452.

66 K. Kansara, A. Mansuri, A. Rajwar, P. Vaswani, R. Singh, A. Kumar and D. Bhatia, Nanoscale Adv, 2023, 5, 2558–2564.

