## Supplementary information for "DNA Tetrahedral Nanocages as a Promising Nanocarrier for Dopamine Delivery in Neurological Disorders"

### Materials and methods

**Materials:** Sigma-Aldrich supplied the following chemicals and reagents: four oligonucleotides with and without Cy3 or Cy5 label, 6× loading dye, 50 bp DNA ladder, mowiol, transferrin-A488, DAPI, and dopamine hydrochloride. Gibco supplied DMEM (Dulbecco's modified Eagle's medium), Medium/F12 (DMEM/F12),), PenStrep, and Trypsin-EDTA (0.25%), while HyClone supplied phosphate buffer saline (PBS). Retinoic acid was purchased from Sigma-Aldrich. Himedia provided ethidium bromide, TEMED, ammonium persulfate, paraformaldehyde, DMSO, adherent cell culture plates, and nuclease-free water. Acrylamide/bis(acrylamide) sol 30% and Tris-Acetate EDTA (TAE) were purchased from GeNei. Magnesium chloride was purchased from SRL, India. All chemicals and reagents were purchased from commercial suppliers and used without further purification unless noted.

**Synthesis, Characterization of DNA-TD and TD-dopamine Complex:**

**Synthesis of self-assembled DNA tetrahedral nanocages:** DNA tetrahedron was synthesized using a one-pot synthesis method. Four single-stranded 55-base oligonucleotides were mixed in equimolar concentration in a reaction solution containing 2 mM MgCl2. The thermal annealing reaction was performed in a PCR instrument with cycling conditions from 95 °C to 4 °C temperature. The reaction mixture was first heated at 95 °C for 30 min and then gradually cooled to 4 °C. It is performed by following these steps: starting temperature of 95°C, decreasing the temperature by 5°C, holding for 15 minutes, and repeating the process until reaching a temperature of 4°C. The T4 oligonucleotide was labeled with Cy3 or Cy5 dye at 5′ ends for imaging purposes.

**Electrophoretic Mobility Shift Assay (EMSA)**: Electrophoretic mobility shift assay (EMSA) was performed using Native-PAGE. 10% polyacrylamide gel was prepared to study the tetrahedral structure formation. The gel was run at 90 V for 90 min.[1] The gel was stained with EtBr stain and visualized using the Gel Documentation system (Biorad ChemiDoc MP Imaging System). The quantification measurement of the migration distance of TD-Dopamine complexes with respect to DNA-TD (100%) was done.

**Dynamic Light Scattering (DLS) and Zeta Potential.** The size-based characterization of DNA tetrahedrons, Dynamic Light Scattering (DLS), and Zeta potential measurement were done with the Malvern analytical Zeta sizer Nano ZS instrument. The data was plotted in Origin Pro software, followed by Gaussian fit.

**Atomic Force Microscopy (AFM).** The morphology-based characterization was performed using Atomic Force Microscopy (AFM).[2,3] Samples were prepared by following established protocols. 5-10 µL aliquots TD and TD-Dopamine complexes were spread over the freshly cleaved mica surface and dried. The imaging was performed with Bruker Nano wizard Sense+ Bio AFM installed at IITGN Gandhinagar Gujrat. Particle size distribution of AFM Images was done by ImageJ imaging software, and the measured data is plotted in Origin pro software.[4,5]

**Fluorescence Spectroscopy:**
All fluorescence titration experiment were recorded by Fluorescence Spectrophotometer Model: Horiba Jobin Yvon Fluorolog-3 at room temperature.

**Circular Dichroism Spectroscopy:** The titration experiment of TD against increasing concentration of Dopamine was performed on the JASCO J-815 CD Spectrometer. All the CD spectra were collected between 190 nm to 400 nm and each spectrum was the average of 3 scans.

**Molecular Dynamics Simulation details:**

The initial structure of the TD was built using the Polygen[6], the Dopamine structure corresponding to the PubChem CID: 681, was extracted from the PubChem database. Then, the geometry optimization of the Dopamine molecule was carried out by using the B3LYP functional together with a 6-311G* basis set with the Gaussian 09 program. The built structures were then solvated in a cubic box with a buffer length of 15 Å using the TIP3P water model[7] with xLEAP module of AMBER20.[8] In this way, all atoms of the solute molecules will be 15 Å apart from the edge of the water box. The negative charges of the phosphate backbones were neutralized by adding the appropriate number of $\mathrm{Mg}^{2+}$ ions. To achieve systems with the desired salt concentration of 0.15 M, we added appropriate numbers of $\mathrm{MgCl_2}$ molecules. Periodic boundary conditions were used in all three dimensions to mimic the bulk properties of the system. All MD simulations were carried out using PMEMD.cuda module of AMBER20. The interactions of TD atoms along with TIP3P waters are defined by leaprc.DNA.bsc1[9,10] forcefield. For, divalent ion ($\mathrm{Mg}^{2+}$), the Li-Merz ion parameters were chosen. The interaction parameters for the protonated dopamine were generated from the general Amber forcefield (GAFF)[11] with the restrained electrostatic potential (RESP)[12] charges. Following that, the energy of the solvated systems was minimized using the steepest descent method for the first 4000 steps, followed by the conjugate gradient method for the next 6000 steps. All of the TD and Dopamine atoms were held fixed by a harmonic potential with a force constant of 500 Kcal.$\mathrm{mol}^{-1}.\mathrm{Å}^{-2}$, so that ions and water molecules can orient themselves from their initial ordered configuration to remove bad contacts. Then, in five successive steps, the positional restrain of the TD atoms were reduced from 500 Kcal.$\mathrm{mol}^{-1}.\mathrm{Å}^{-2}$ to 5 Kcal.$\mathrm{mol}^{-1}.\mathrm{Å}^{-2}$, leading to their eventual equilibration with no constraints.

The systems were heated slowly in four steps: first, from 10 to 50 K for 6,000 MD steps, secondly, from 50 to 100 K for 12,000 steps, from 100 to 200 K for 10,000 steps, from 200 to 300 K for 10,000 steps, and finally, at 300 K for another 12000 MD steps. Positional restraints were applied to the heavy atoms of the solute during heating with a force constant of 20 Kcal.$\mathrm{mol}^{-1}.\mathrm{Å}^{-2}$. Following heating, the systems undergo a 5 ns NPT equilibration with a Berendsen weak coupling method to adjust the box size with a target pressure of 1 bar.[13,14] All bonds involving hydrogens were constrained using the Shake algorithm. This allowed us to use an integration time step of 2 fs. The electrostatic interactions were treated using the Particle Mesh Ewald method with a cutoff of 10 Å.[15] Following the equilibration, the systems were simulated for production run using a Langevin thermostat with a coupling constant of 1 ps for 200 ns at a temperature of 300 K.[16] Similar simulation methodologies have been implemented in some of the previous studies.[17,18]

All the simulation trajectories were analyzed with the Visual Molecular Dynamics (VMD), CPPTRAJ for modelling and the analysis of molecular systems. Xmgrace tool (http://plasma-gate.weizmann.ac.il/Grace/)), Pymol and Python Matplotlib were also used to visualize the data and construction of the figures shown.[19-21]

**Cytotoxicity effects of TD-Dopamine complexes with MTT assay:**[22] MTT assay was performed to assess the effect of cytotoxicity of TD-Dopamine complexes and dopamine alone. SH-SY5Y cells were seeded in 96 well plates at the seeding density of 10,000 cells per well. The culture plates were incubated at 37˚C for 24 h. The cells were treated with different ratios of Td, dopamine and TD-dopamine complexes. The untreated cells served as a control. After incubation, 0.5mg/ml of 3-(4,5-Dimethylthiazol-2-yl)-2,5-diphenyltetrazolium Bromide (MTT) solution was added to each well and incubated at 37˚C for 4 h. The solution was removed, replaced with dimethyl sulfoxide (DMSO) in each well, and incubated in the dark for 15 minutes to dissolve the formazan crystal. The multiwell microplate reader was used to measure absorbance at 570 nm. The cell viability percentage was calculated using the formula:

$$Cell\ viability\ (\%) = \frac{Absorbance\ of\ the\ sample}{Absorbance\ of\ the\ control}\ \mathrm{x}\ 100$$

**Differentiation of SH-SY5Y cells:** The differentiation experiment was done by following a well-established protocol.[23–25] Differentiation of human neuroblastoma SH-SY5Y cells was performed as explained in the earlier studies. Briefly, on day 0, undifferentiated SH-SY5Y at 60–80% confluency was trypsinized (0.25% trypsin), and 5 × 105 cells were seeded in a T-25 flask. Around $10^5$ cells per well were seeded on a 10mm glass coverslip in a four-well plate 24h before the experiment in regular culture media. On the first day, cells were treated with DNA-TD, dopamine TD-dopamine complex at different differentiation media (DM)-1 composition. After the treatment, the cells were incubated at 37oC. The same treatment was repeated on day 3 and day 5. On day 7 and day 9 the same treatment was done in DM-2, and on day 11 was treated in DM-3. The compositions of differentiation media (DM) are: **DM-1:** For 10 mL of media, 9.73mL of (DMEM/F12 (1:1) with 1x Pen/Strep) + 250µL of 2.5% FBS + 20µL of RA 5mM; **DM-2:** For 10 ml of media (9.88mL of (DMEM/F12 (1:1) with 1x Pen/Strep + 100µL of 2.5% FBS + 20µL of RA 5mM; **DM-3:** For 10 ml of media Neurobasal 200µL of 1M KCl + 100µL of 100X Glutamaxl + 200µL B-27 + 20µL of RA 5mM. The same procedure was followed to check the differentiation ability of DNA-TD and TD-Dopamine complex testing samples replaced the RA. Untreated cells were cultured only in DMEM F12 complete media and used as a control. Retinoic acid was used as a standard differentiator.

**Cellular Uptake Assay:** The cellular uptake of TD and TD-dopamine complexes into differentiated and undifferentiated SH-SY5Y cells was examined. Approximately $10^5$ per well cell counts were seeded overnight on a glass coverslip in a 4-well plate. Before treatment, the seeded cells were washed with 1X PBS buffer thrice and then incubated in serum-free media for 30 min at 37 °C with 5% $CO_2$ in a humidified incubator. After the wash, the cells were treated with DNA-TD and TD-Dopamine complexes in different compositions to assess their cellular internalization. The treated cells were fixed for 15 minutes at 37 °C with 4% paraformaldehyde and rinsed three times with 1X-PBS. The cells were then permeabilized with 0.1% Triton-X100 and stained with 0.1% phalloidin to visualize the actin filaments. Then the cells were washed thrice with 1X PBS and mounted onto a glass slide using mowiol containing Hoechst to mark the nucleus.

**Zebrafish maintenance**

The zebrafish used in this study was of the Assam wild-type strain and was grown in the lab from embryo to adult stage. The husbandry and maintenance of zebrafish were executed according to our previously published protocol.[26] Briefly, the lab conditions were maintained according to the laboratory conditions explained on ZFIN (Zebrafish Information Network), including a 14 h light / 10 h dark cycle at a temperature of 26 – 28°C. The fish were kept in a 20 L tank with aeration pumps. The water of the tanks was prepared by adding 60 mg/L sea salt (Red Sea Coral Pro salt). The various parameters were maintained to mimic the natural environment i.e., pH (7-7.4), conductivity (250 – 350 µS), TDS (220– 320 mg/L), salinity (210–310 mg/L), and dissolved oxygen (> 6 mg/L) using a multi-parameter instrument (Model PCD 650, Eutech, India). The zebrafish were fed brine shrimp (live artemia) and basic flakes (Aquafin). They were fed the artemia twice and flakes once a day. The breeding setup was prepared in the lab with a ratio of 3 females and 2 males in the breeding chamber. Post breeding, the embryos were collected into E3 medium in a sterile petri dish and kept in a BOD incubator (MIR -154, Panasonic, Japan) at 28.0°C. The embryos were raised in the same medium for three days and then the healthy larvae were used for the experiment.

**The survival rate, malformations and heart rate analysis**

The survival rate, malformations and heart rate studies were conducted by distributing 15 zebrafish larvae per well in a six-well plate (Corning, NY, USA). Volume in each well was made up to 5mL. One well was designated as a control in each group without any nanostructure's exposure. The final concentration of TD was 300 nM and TD-Dopamine was in a 1:100 ratio (Dopamine at concentrations 100 eq.,). Our set-up consists of a stereo-zoom microscope utilized to measure the heartbeats of zebrafish larvae manually. Heart rate studies were conducted by distributing 15 larvae per well in a six-well plate (Corning, NY, USA). Volume in each well was made up to 5mL. One well was designated as a control in each group without any nanostructure's exposure. The final concentration of TD was 300 nM and TD-Dopamine was in a 1:100 ratio.

***In vivo* uptake of TD-Dopamine in Zebrafish larvae**

*In vivo*, uptake assays were performed according to the Organization for Economic Cooperation and Development (OECD) guidelines. At 72 h after hatching, corresponding to 8 days post-fertilization (dpf), the dead embryo was removed and the remaining larvae were placed in six-well plates (Corning, NY, USA) with 15 zebrafish larvae in each well. Three groups of zebrafish larvae were treated with TD, Dopamine and TD-Dopamine at concentrations of 300nM and 1:100 (TD: Dopamine) ratio and incubated for 4 hours. One well was designated as a control in each group without nanoparticles. Post-treatment, the medium was replaced with fresh E3 media, and zebrafish larvae were washed to remove the excess TD and TD-Dopamine and fixed with a fixative solution (4% PFA) for 2 minutes. Post fixation, the zebrafish larvae were mounted with mounting solution Mowiol and allowed to dry for further confocal imaging analysis.

**Confocal Microscopy:** Leica TCS SP8, Confocal Scanning Laser Microscope was used for imaging the fixed cells and fixed embryos. Different lasers were used to activate fluorophores, 405 nm for Hoechst, 488 nm for phalloidin, 561 nm for TD-Cy3 and 633 nm for Cy5. Further, the Image analysis was done by Fiji ImageJ software for image analysis. The background signal (error) was subtracted and the intensity of each cell was calculated. The intensity of 40−55 cells was measured for the quantification of cellular uptake experiments. All qualified data is plotted in Graphpad Prism software.[1,27]

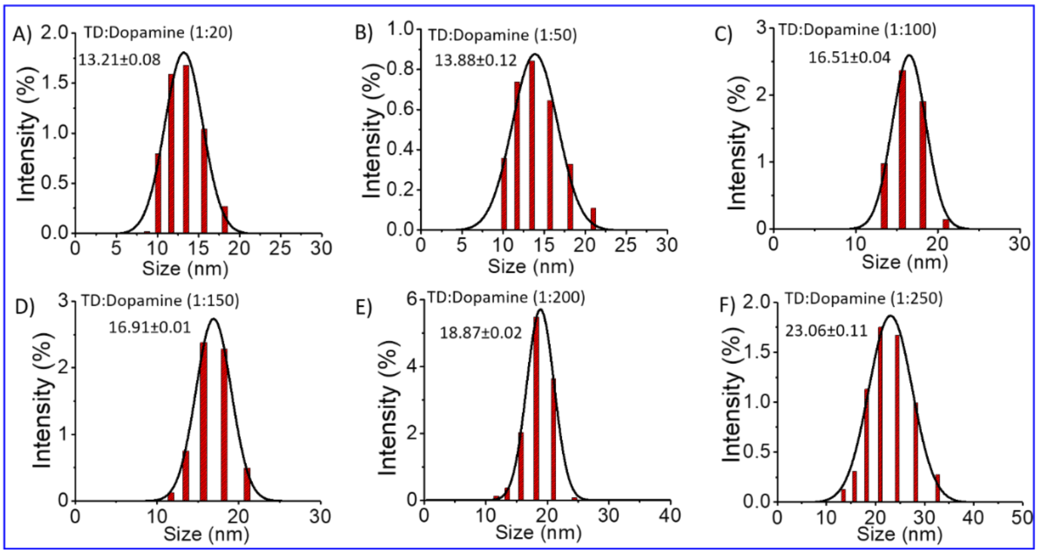


**Figure S1:** DLS bar graph of TD-dopamine nanoparticles in different TD:dopamine compositions. A) 1:20, B) 1:50, C) 1:100, D) 1:150, E) 1:200 and F) 1:250.

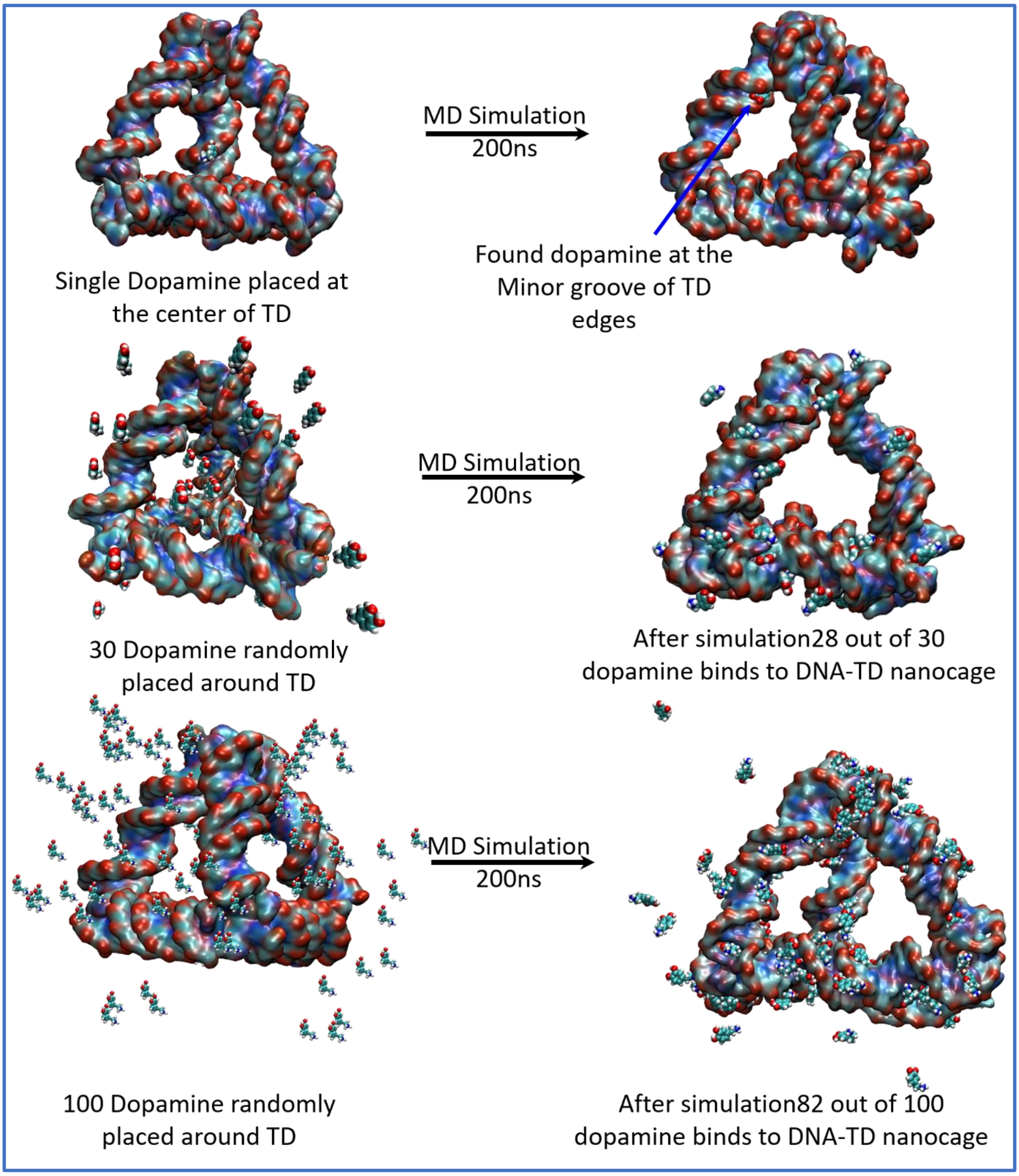


**Figure S2:** depicts the configuration of dopamine molecules around DNA TD before and after MD simulation for 200ns.

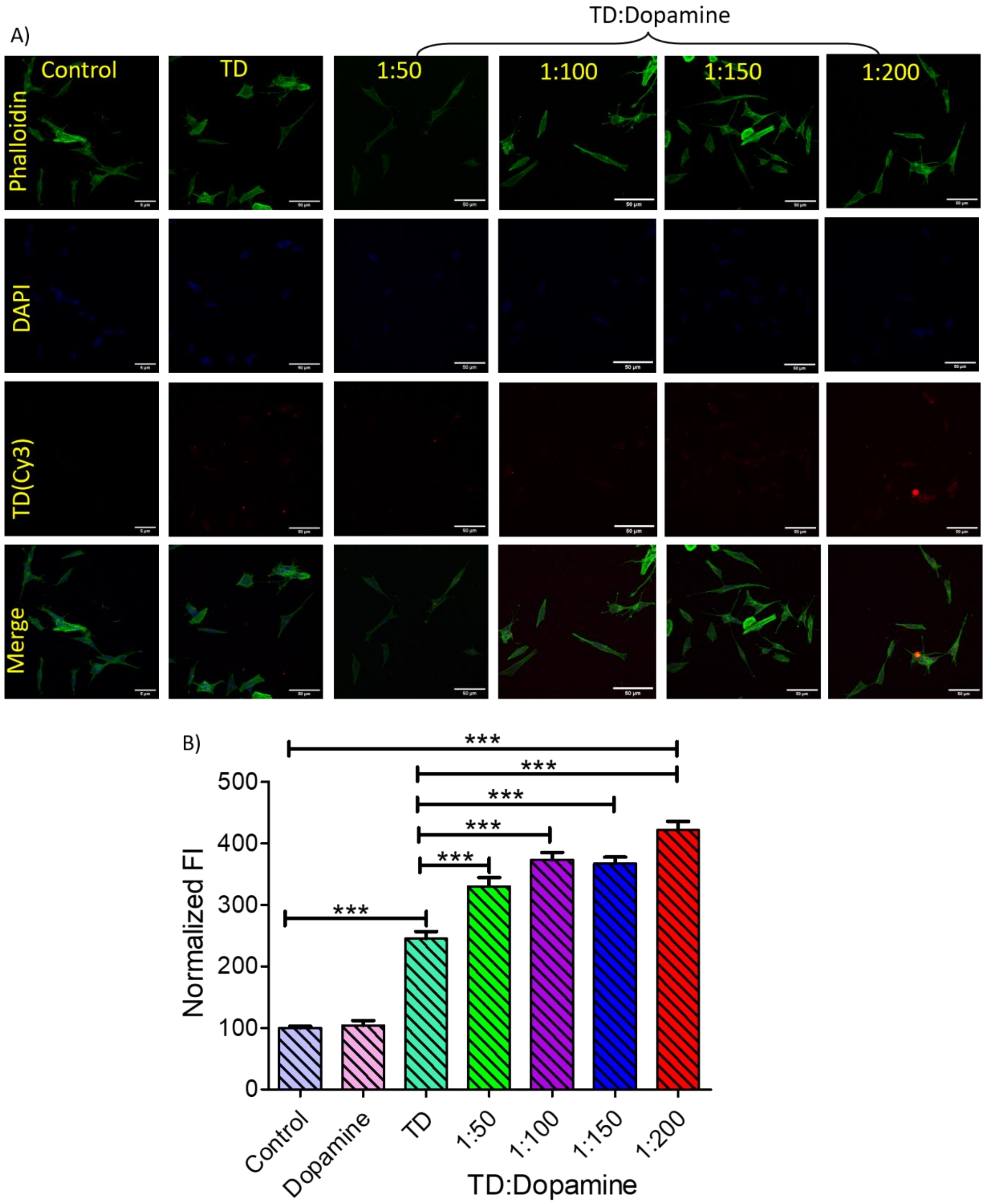


**Figure S3:** Cellular uptake of TD-Dopamine complex nanostructures into undifferentiated SH-SY5Y neuroblastoma cell lines. A) Confocal imaging of differentiated SH-SY5Y neuroblastoma cells treated with DNA Cy3 TD (150 nm) and with Cy3 TD (150 nm)–Dopamine in different ratios of TD and Dopamine for 15 min. The green channel represents an actin cytoskeleton stained with Phalloidin-A488, the blue channel represents nuclei stained with DAPI, the red channel represents TD(Cy3) uptake, and the lower panel represents merged images. The scale bar is 50µm. B) Quantifying TD–Cy3 uptake in differentiated SHSY-5Y neuroblastoma cells from panel A. D) and E) Bar graphs representing cell viability (%) obtained from MTT assay test against different concentrations of Dopamine with TD (150nm) and without TD. ***Statistically significant p-value ($p < 0.0001$).

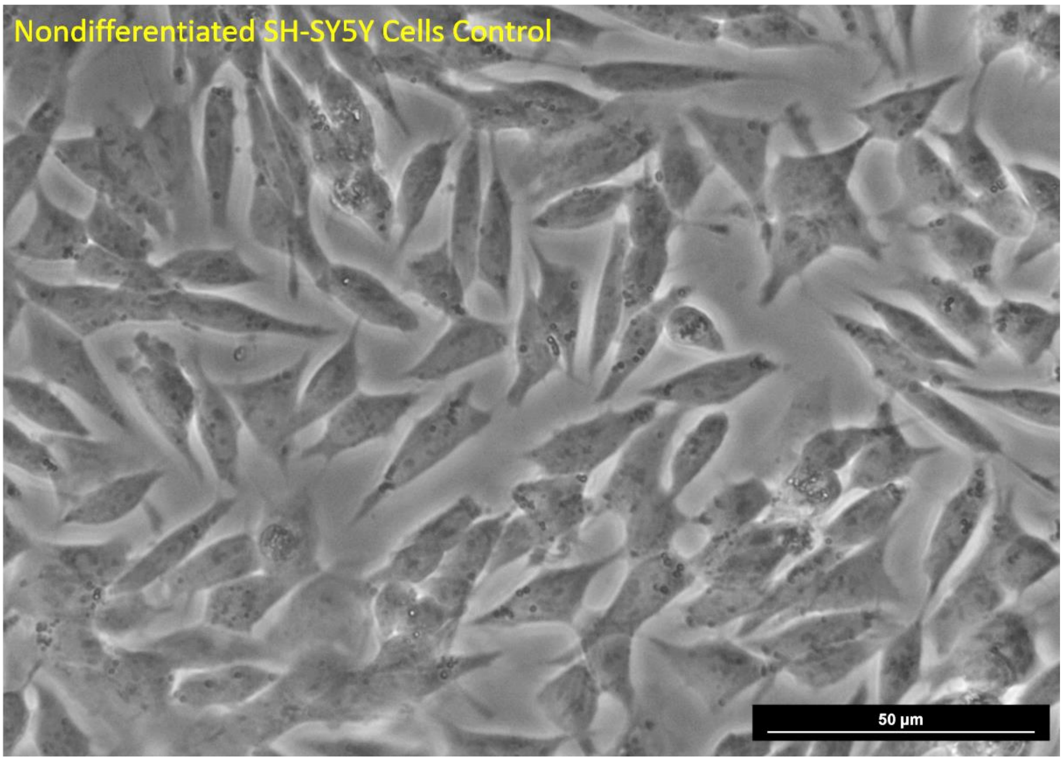


**Figure S4:** Optical image of non-differentiated SH-SY5Y cells

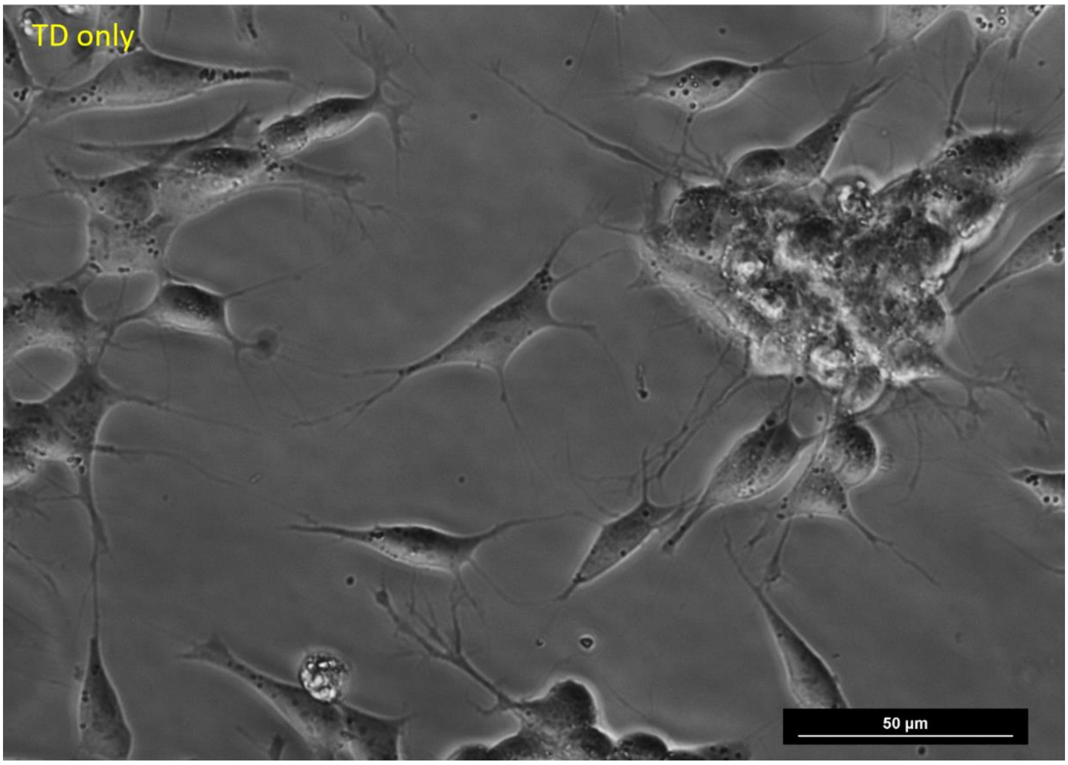


**Figure S5:** Optical image of SH-SY5Y cells differentiated by DNA-TD (150nM)

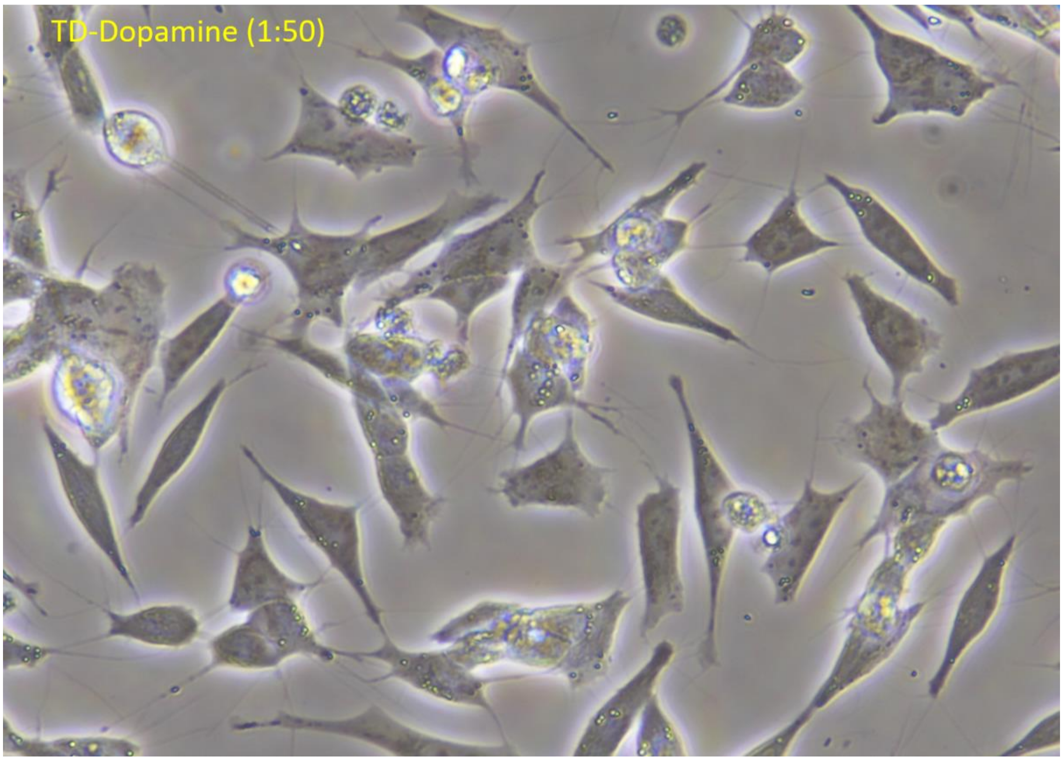


**Figure S5:** Optical image of SH-SY5Y cells differentiated by TD-dopamine complex in composition of (1:50)

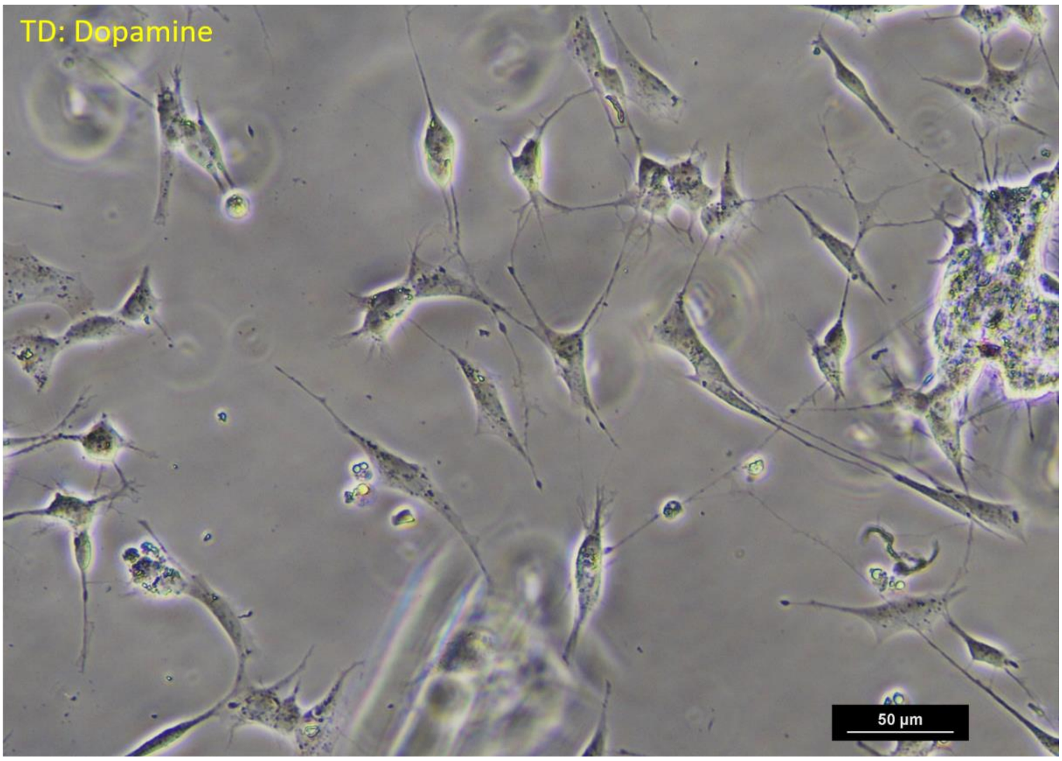


**Figure S6:** Optical image of SH-SY5Y cells differentiated by TD-dopamine complex in composition of (1:200)

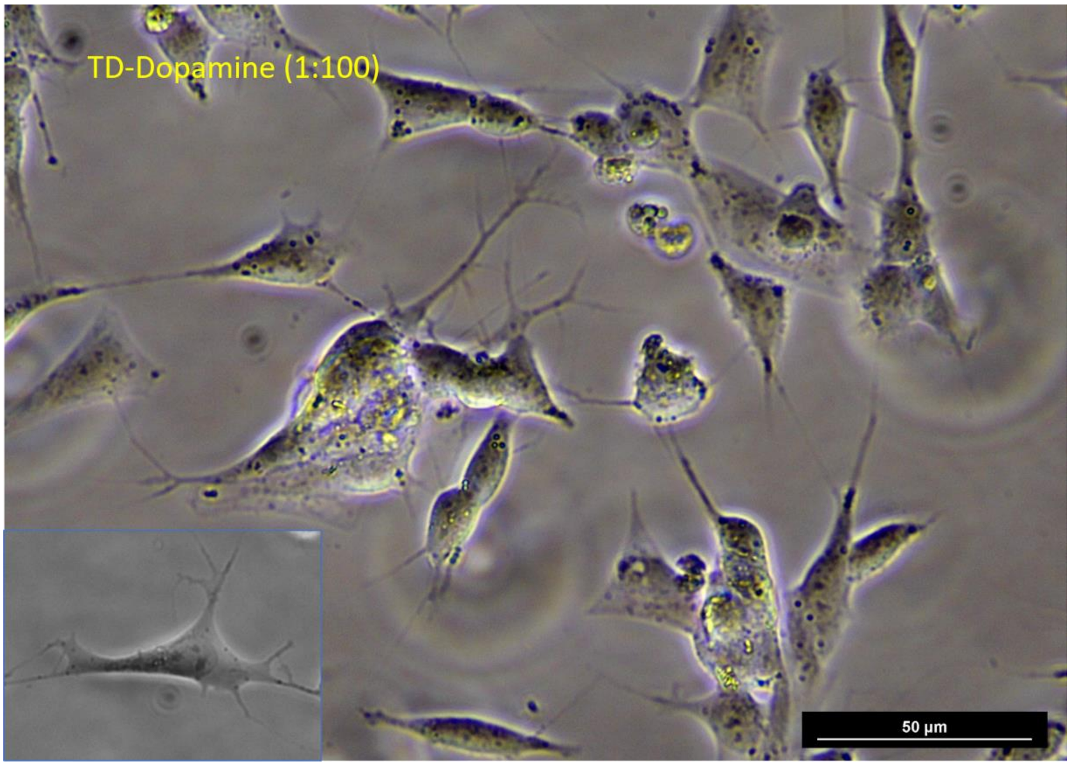


**Figure S7:** Optical image of SH-SY5Y cells differentiated by TD-dopamine complex in composition of (1:100)

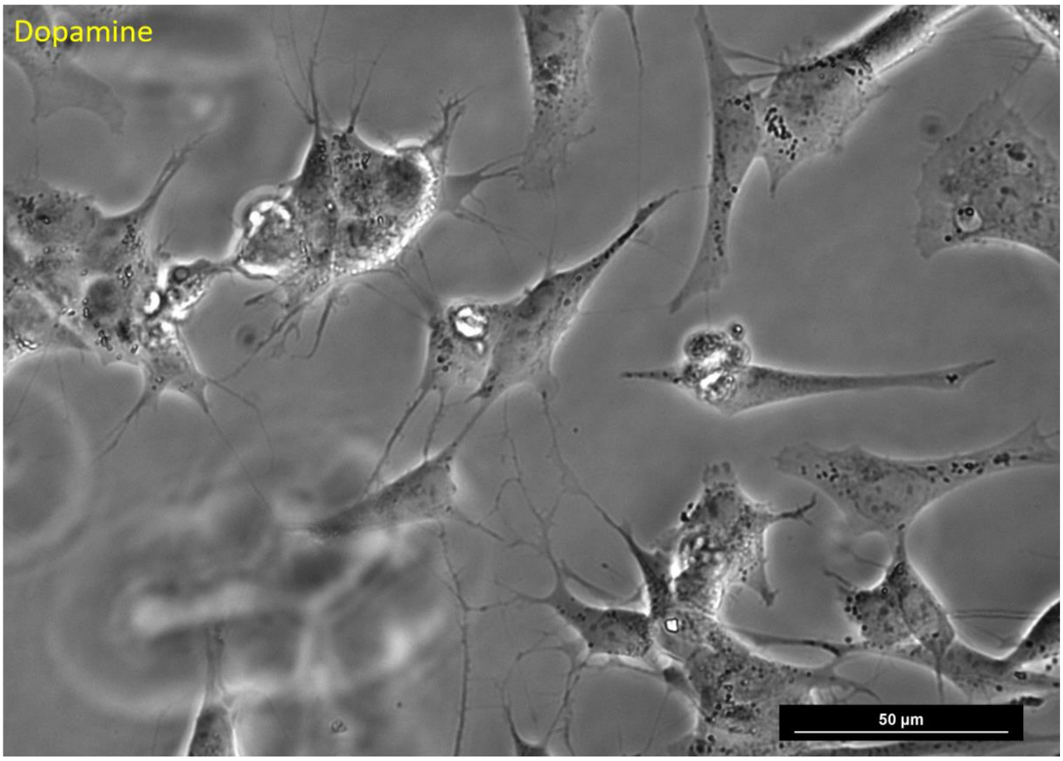


**Figure S8:** Optical image of SH-SY5Y cells differentiated by Dopamine (30μM).

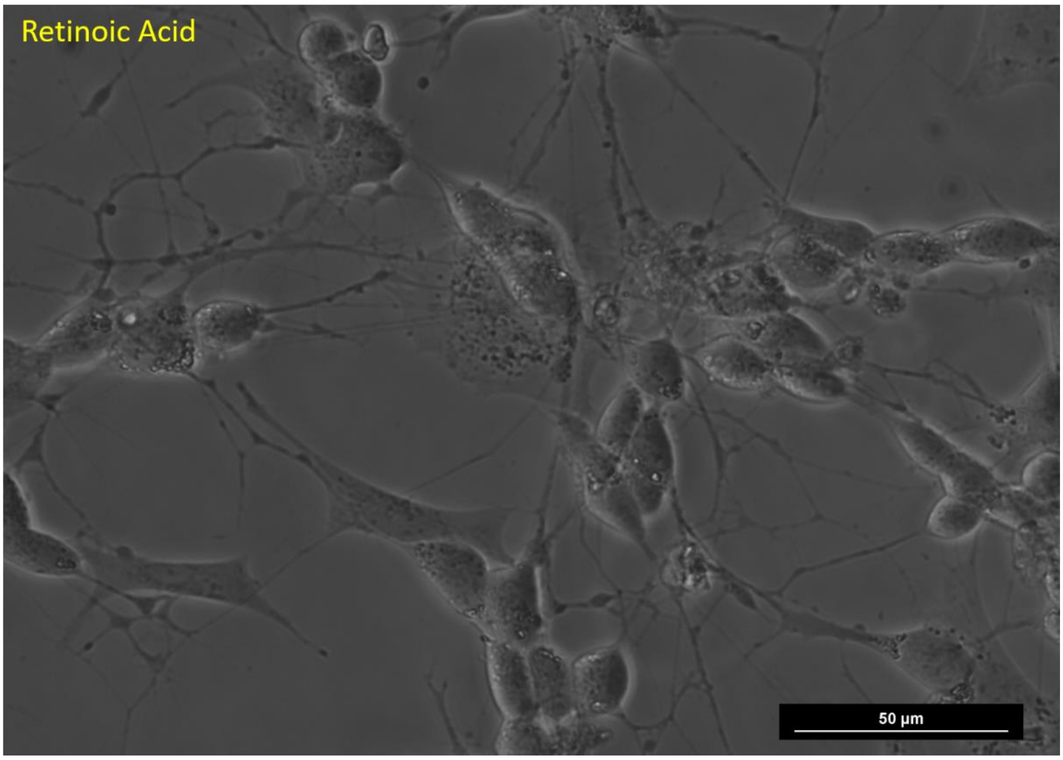


**Figure S9:** Optical image of SH-SY5Y cells differentiated by Retinoic Acid.

**References:**


1 A. Rajwar, S. R. Shetty, P. Vaswani, V. Morya, A. Barai, S. Sen, M. Sonawane and D. Bhatia, *ACS Nano*, 2022, **16**, 10496–10508.

2 R. Singh, N. Kumar Mishra, V. Kumar, V. Vinayak and K. Ballabh Joshi, *ChemBioChem*, 2018, **19**, 1630–1637.

3 R. Singh, N. K. Mishra, P. Gupta and K. B. Joshi, *Chem Asian J*, 2020, **15**, 531–539.

4 N. Singh, S. Sharma, R. Singh, S. Rajput, N. Chattopadhyay, D. Tewari, K. B. Joshi and S. Verma, *Chem Sci*, 2021, **12**, 16085–16091.

5 N. Singh, R. Singh, S. Sharma, K. Kesharwani, K. B. Joshi and S. Verma, *New Journal of Chemistry*, 2021, **45**, 153–161.

6 C. Alves, F. Iacovelli, M. Falconi, F. Cardamone, B. Morozzo della Rocca, C. L. P. de Oliveira and A. Desideri, *J Chem Inf Model*, 2016, **56**, 941–949.

7 W. L. Jorgensen, J. Chandrasekhar, J. D. Madura, R. W. Impey and M. L. Klein, *J Chem Phys*, 1983, **79**, 926–935.

8 Case, David A., et al. "Amber 2020: University of california." *San Francisco* (2020).

9 T. E. Cheatham and D. A. Case, *Biopolymers*, 2013, n/a-n/a.

10 I. Ivani, P. D. Dans, A. Noy, A. Pérez, I. Faustino, A. Hospital, J. Walther, P. Andrio, R. Goñi, A. Balaceanu, G. Portella, F. Battistini, J. L. Gelpí, C. González, M. Vendruscolo, C. A. Laughton, S. A. Harris, D. A. Case and M. Orozco, *Nat Methods*, 2016, **13**, 55–58.

11 J. Wang, R. M. Wolf, J. W. Caldwell, P. A. Kollman and D. A. Case, *J Comput Chem*, 2004, **25**, 1157–1174.

12 C. I. Bayly, P. Cieplak, W. Cornell and P. A. Kollman, *J Phys Chem*, 1993, **97**, 10269–10280.

13 H. J. C. Berendsen, J. P. M. Postma, W. F. van Gunsteren, A. DiNola and J. R. Haak, *J Chem Phys*, 1984, **81**, 3684–3690.

14 Van Gunsteren, W. F.; Berendsen, H. J. A leap-frog algorithm for stochastic dynamics.Molecular Simulation 1988, 1, 173–185.

15 T. Darden, D. York and L. Pedersen, *J Chem Phys*, 1993, **98**, 10089–10092.

16 Davidchack, R. L.; Handel, R.; Tretyakov, M. Langevin thermostat for rigid body dynamics. The Journal of chemical physics 2009, 130, 234101.

17 H. Joshi, A. Kaushik, N. C. Seeman and P. K. Maiti, *ACS Nano*, 2016, **10**, 7780–7791.

18 P. Kumbhakar, I. Das Jana, S. Basu, S. Mandal, S. Banerjee, S. Roy, C. C. Gowda, A. Chakraborty, A. Pramanik, P. Lahiri, B. Lahiri, A. Chandra, P. Kumbhakar, A. Mondal, P. K. Maiti and C. S. Tiwary, *Physical Chemistry Chemical Physics*, 2023, **25**, 17143–17153.

19 W. Humphrey, A. Dalke and K. Schulten, *J Mol Graph*, 1996, **14**, 33–38.

20 D. R. Roe and T. E. Cheatham, *J Chem Theory Comput*, 2013, **9**, 3084–3095.

21 J. D. Hunter, *Comput Sci Eng*, 2007, **9**, 90–95.

22 U. Singh, A. G. Teja, S. Walia, P. Vaswani, S. Dalvi and D. Bhatia, *Nanoscale Adv*, 2022, **4**, 1375–1386.

23 M. M. Shipley, C. A. Mangold and M. L. Szpara, *Journal of Visualized Experiments*, , DOI:10.3791/53193.

24 J. Kovalevich and D. Langford, 2013, pp. 9–21.

25 P. Hivare, A. Gangrade, G. Swarup, K. Bhavsar, A. Singh, R. Gupta, P. Thareja, S. Gupta and D. Bhatia, *Nanoscale*, 2022, **14**, 8611–8620.

26 K. Kansara, A. Paruthi, S. K. Misra, A. S. Karakoti and A. Kumar, *Environmental Pollution*, 2019, **255**, 113313.

27 U. Singh, A. G. Teja, S. Walia, P. Vaswani, S. Dalvi and D. Bhatia, *Nanoscale Adv*, 2022, **4**, 1375–1386.
